# Endogenous retrovirus activation disrupts postsynaptic organization in human glutamatergic neurons

**DOI:** 10.64898/2026.10.21.760639

**Authors:** Carrie Davis-Hansson, Raquel Garza, Claire Piochon, Jenny G. Johansson, Volker Busskamp, Johan Jakobsson, Emily M. Johansson

## Abstract

Endogenous retroviruses (ERVs) comprise a substantial fraction of the human genome, yet whether their expression can directly alter human neuronal function remains unknown. Because glutamatergic signaling is fundamental to brain function and frequently disrupted in brain disorders, we asked whether activation of the primate-specific ERV-W group, associated with neurological disease, perturbs human glutamatergic neurons. We used CRISPR activation of the LTR17-ERV-W consensus sequence to examine its effects on neuronal and synaptic function, inducing transcription from multiple proviral loci, including five with intact open reading frames. Functionally, ERV-W activation selectively altered the postsynaptic compartment, reducing PSD-95 and NMDA receptor abundance. These changes were accompanied by altered excitatory synaptic transmission and reduced intrinsic excitability, whereas activation of ERVW-1/Syncytin-1 alone did not reproduce the phenotype. Our findings identify ERV-W activation as a modulator of human neuronal function with potential relevance to ERV-associated brain disorders.

## Introduction

Glutamatergic synapses mediate the main excitatory signaling in the human brain and are central to complex cognitive functions^1,2^. These synapses mediate fast excitatory neurotransmission through ionotropic receptors, such as alpha-amino-3-hydroxy-5-methyl-4-isooxazole-propionic acid-(AMPA) and N-methyl-d-aspartate-(NMDA) receptors and play a key role in synaptic plasticity that shape neuronal networks^3^. Synaptic integrity is tightly regulated by coordinated pre- and postsynaptic molecular mechanisms that fine-tune synaptic strength and maintain network stability^4–6^. Consequently, even subtle perturbations in synaptic composition can propagate across neural circuits and lead to measurable deficits in information processing and cognitive performance. Accordingly, disturbance of glutamatergic synapses is a hallmark of several neuropsychiatric and neurodegenerative disorders where disrupted activity and integrity often precede neuronal loss and correlates with loss of cognitive functions^5,7–10^. Despite this, the molecular mechanisms underlying early synaptic impairment in human brain disorders remain poorly understood.

There is a growing body of evidence linking transposable elements (TE), that comprise at least 50% of our genome^11,12^, to human brain disorders^13–17^. For instance, increased TE expression has been found to correlate with senescence and other age-related cellular phenotypes^18,19^, and has been observed in both neurodevelopmental, neuropsychiatric and neurodegenerative disorders^17,20–25^. Such aberrant TE expression is likely to have multiple functional consequences in the brain. For example, it is well established that TEs can act as potent cis-regulatory elements in human neural cells, for example by acting as alternative promoters or enhancers^17,26–29^. Thus, disease-related changes in TE activity can potentially reshape gene regulatory networks. In addition, we and others have recently demonstrated that TE activation in the adult brain elicits a neuroinflammatory response in rodent models^30^ and, that TE expression correlates with an innate immune response in both human neurodegenerative and neuropsychiatric states^17,31,32^.

Human retrotransposons are divided into two main categories, Long Terminal Repeat (LTR) retrotransposons and non-LTR retrotransposons, based on their structure and mechanism of transposition. Similar to LTR retrotransposons, endogenous retroviruses (ERVs) that derive from heritable retrovirus infections in the germline and now make up about 8% of our genome ^33^, also have LTRs flanking their proviral genes (*gag, pro, pol* and *env*). Although most ERV loci occur as solitary LTRs resulting from homologous recombination between the two proviral LTRs, many ERVs have retained their genes and even protein-coding potential^33–35^. ERV expression in the human brain is widespread but tightly regulated, varying across neuronal populations, brain regions, and developmental stages^17,36,37^. However, the functional significance of this expression remains poorly understood, particularly which individual ERV elements are active in specific neuronal populations and how they influence neuronal function.

There is accumulating evidence that implicates ERVs as potential contributors to brain disorders. For example, increased levels of transcripts and peptides from ERV-W, a largely primate specific group of ERVs have been found in several neurological and psychiatric disorders, including Multiple sclerosis, Parkinson’s disease, schizophrenia, and autism^17,21,31,38,39^. Furthermore, studies in rodent models have linked ERV-W expression to pathological processes in the mammalian brain, demonstrating that ERV-W activation can have a significant impact on synaptic plasticity and cognitive function^32,40^. However, the underlying mechanism by which ERV-W expression affects cell communication and synaptic transmission in human neurons is unclear, yet this is a fundamental question with potential clinical implications. Here, we developed a CRISPR-activation system for efficient and specific induction of ERV-W transcription in human cells. Activation of endogenous ERV-W loci in human neurons disrupted postsynaptic organization and neuronal excitability, resulting in a destabilized synaptic state and dysregulated postsynaptic signaling. Together, these findings demonstrate that ERV-W expression impairs excitatory postsynaptic integrity in human neurons.

## Results

### ERV expression is silenced during human neuronal maturation

The human genome is estimated to harbor around 3,677 ERV loci based on RetroTector predictions with scores >300, reflecting the extent to which sequences match known retroviral motifs and structural features^41^. These loci correspond to approximately 2865 proviral ERV predictions (at least one LTR region). There are also numerous ERV-derived solo LTRs in the human genome (Figure 1A)^16,41^. To investigate ERV expression in mature neurons, we optimized a forward reprogramming protocol to obtain excitatory neurons from human induced pluripotent stem cells (hiPSCs). This approach directly converts hiPSCs into induced neurons, through the overexpression of the neuronal transcription factors, neurogenin (Neurog)-1 and - 2 (called iNGN neurons), bypassing an intermediate pluripotent or progenitor state^42^. The resulting reprogramming rapidly redirects the transcriptional program of hiPSCs toward a neuronal fate, resulting in the generation of a homogenous population of excitatory neuronal cells within 4 days post induction (dpi) as monitored with ICC and RNA-sequencing (Figure 1B and S1A). We subsequently followed neuronal maturation over time. At 60 dpi the resulting excitatory iNGNs have complex dendritic branches with a high density of synapses as observed by MAP2 and post synaptic density-95 (PSD-95) ICC stainings and spontaneous action potential (AP) activity (Figure 1C), consistent with previous studies^43^. Using GFP-transduced live imaging of iNGNs and, in parallel, high-resolution imaging of fixed neuronal synapses, we could also confirm the presence of well-developed synaptic protrusions at 60 dpi (Figure S1B, C). We traced the morphometric maturation of iNGNs with focus on synapses at 7, 14, 34, 56 and 72dpi. In general, ICC staining against PSD-95 was accompanied by juxtaposed and overlapping labelling of the presynaptic marker vesicular glutamate transporter 1 (vGlut1), which also revealed a nearly complete overlap with another pre-synaptic marker, Synaptophysin (Figure S1D, E). We observed a gradual increase in the PSD-95 density index (number and area of puncta over dendritic length) and number of vGlut1 puncta per neurite over time (Figure S1F, G), with progressively more stable synapses (vGlut1 positive post-synapses) (Figure S1H). Thus, this protocol offers a homogenous model system to study human neuronal function and synaptic maturation.

**Figure 1.**
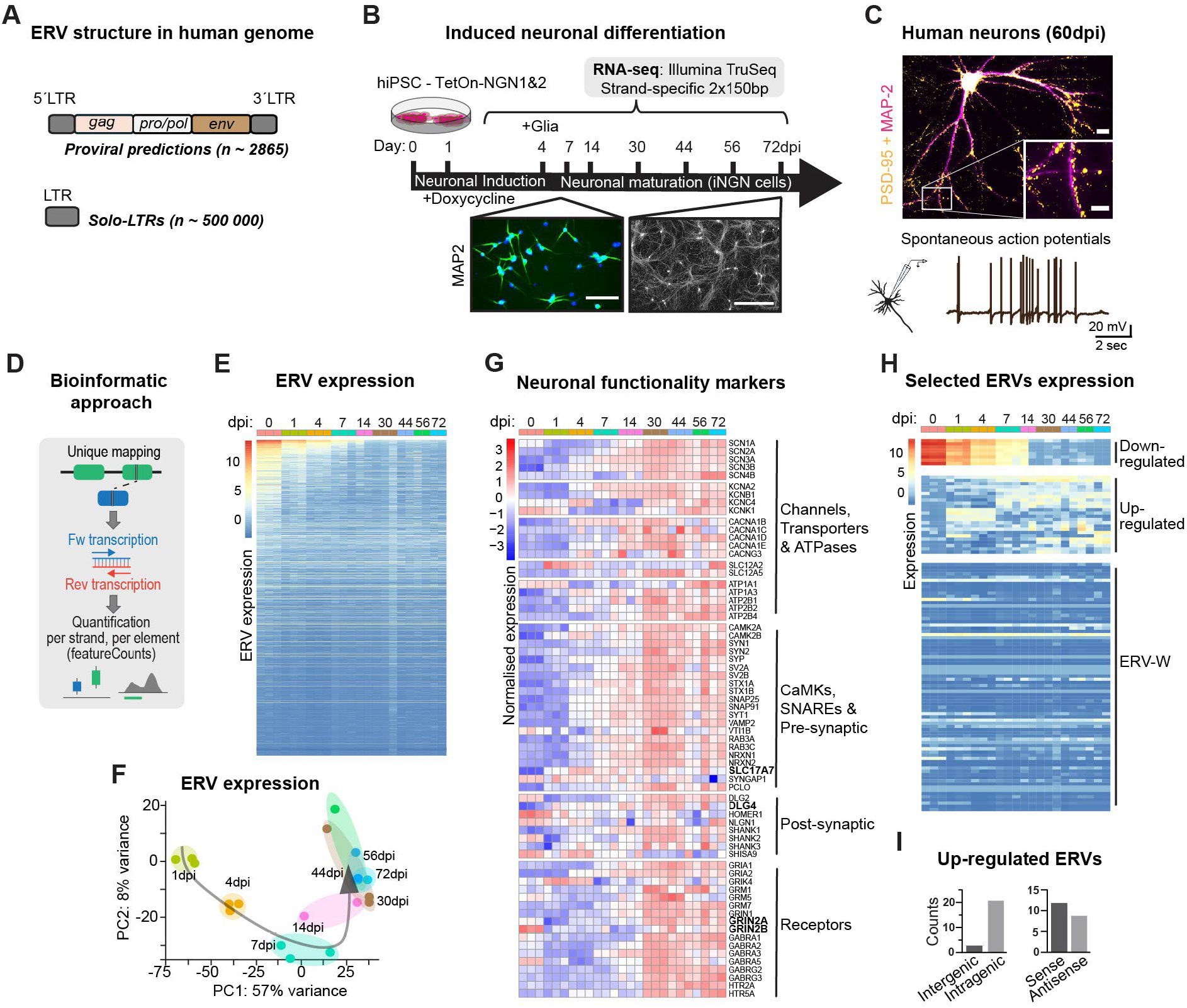
ERV expression in the human genome and in hiPSC derived excitatory neurons. (A) Structure of ERV elements with long terminal repeats (LTRs) (grey boxes) flanking retroviral genes (*gag, pro, pol* and *env*). Around 2865 proviral ERV retrotector predictions with score >300 are detected out of 3677 ERV predictions. (B) Schematic of induced neuronal differentiation. Doxycycline-induced Neurogenin-1/2 overexpression in hiPSCs (iNGNs) generated postmitotic neurons within 4 days post induction (dpi), followed by dissociation and astrocyte co-culture. Samples were collected for bulk RNA sequencing throughout maturation. Scale bars, 100 µm. (C) Representative images of mature human iNGNs at 60 dpi showing PSD-95/MAP2 immunostaining (top) and spontaneous whole-cell patch-clamp recordings (bottom). Scale bars, 20 µm. (D) Bioinformatic rationale to quantify ERV expression by using a unique mapping approach for stranded expression of unique ERV loci. Bulk RNA-sequencing across neuronal differentiation (0, 1, 4, 7, 14, 30, 44, 56, and 72 dpi; n = 2–3 technical (0-7 & 30dpi)/ biological (14-72 dpi) replicates). (E) Heat map showing generally low ERV expression based on the 2,000 most variable uniquely mapped ERV loci (of 3,677 total). (F) PCA of the 500 most variable ERV loci across differentiation/maturation. (G) Heat map of normalized expression of neuronal functional markers, including key pre- and postsynaptic genes and glutamatergic receptors. (H) Representative downregulated ERV-H loci, and upregulated ERV loci during differentiation/maturation, lower panel shows expression of ERV-W proviral predictions (n ∼ 78). (I) Genomic annotation of the upregulated ERVs shown in (H), demonstrating predominantly intragenic localization (21/24 loci) and mainly sense orientation relative to the host gene (12/21 loci), based on RepeatMasker and GENCODE annotations.

To investigate the expression of ERVs during neuronal differentiation and maturation we collected RNA from iNGNs at 0, 1, 4, 7, 14, 30, 44, 56 and 72dpi (Figure 1B). We used an in-house 2 × 150–base pair (bp), polyadenylate [poly(A)]-enriched stranded library preparation for bulk RNA-seq using a reduced fragmentation step to optimize library insert size for TE analysis^29^. Due to different amplification waves of ERV sequences in the genome, it is known that a proportion of ERV reads map ambiguously to the reference genome^33,34^. To avoid potential artefacts resulting from multimapping reads, we therefore used a strand-specific unique mapping approach to analyze the expression of individual ERV loci (Figure 1D).

We found that many proviral ERV loci (∼ 480 with >10 raw counts) are expressed in the pluripotent stage, but this expression was gradually silenced as the neurons mature, leading to an almost complete silencing of ERVs after 30 dpi (Figure 1E). Unsupervised hierarchical clustering of ERV expression revealed that early (1-4dpi) iNGNs group by dpi and separate from later timepoints (30-72dpi) (Figure S1I). Likewise, principal component analysis (PCA) showed that PC1 and PC2 together accounted for 65% of the total variance and organized samples along a maturation trajectory, further indicating that neuronal maturation is the dominant driver of ERV-transcriptional variation in our samples (Figure 1F). The decrease of ERV expression correlated with gene expression onset of genes related to neuronal functionality, such as channels and transporters, pre- and postsynaptic genes (e.g. *SCL17A7* and *DLG4*) including receptors for neurotransmitters and glutamate receptor subunits that are related to synaptic maturation (*GRIN2B* and *GRIN2A*) (Figure 1G).

The ERVs that were highly expressed at the pluripotent stage and subsequently downregulated upon neuronal maturation primarily belonged to the ERV-H group, consistent with previous reports describing high ERV-H expression in human pluripotent cells^44^. While most proviral ERV-W predictions remained lowly transcribed across all time points, a small number of ERVs from diverse ERV groups were upregulated during neuronal maturation (Figure 1H). However, most of the upregulated ERVs were located within genes and transcribed in the same orientation as their host coding genes (Figure 1I), suggesting that their expression largely reflects passive co-transcription as part of host-gene transcripts rather than independent ERV transcription. In conclusion, the vast majority of ERVs, including ERV-W, remained transcriptionally silent across the different stages of neuronal maturation, demonstrating that ERV elements are largely silenced as human neurons mature.

### CRISPRa-mediated activation of endogenous ERV-W proviruses in human iPSCs

To investigate the consequences of increased ERV expression during neuronal maturation, we focused on the ERV-W group based on accumulating evidence linking aberrant ERV-W expression to several neurological and psychiatric disorders^13,15,17,32,45^. We identified 78 predicted LTR17-ERV-W proviruses (hereafter referred to as ERV-W), with LTR17 representing the LTR most frequently flanking ERV-W proviral sequences^46^, and 1100 solo-LTR17s, which were equally distributed between intra- and intergenic regions (Figure 2A,B).

**Figure 2.**
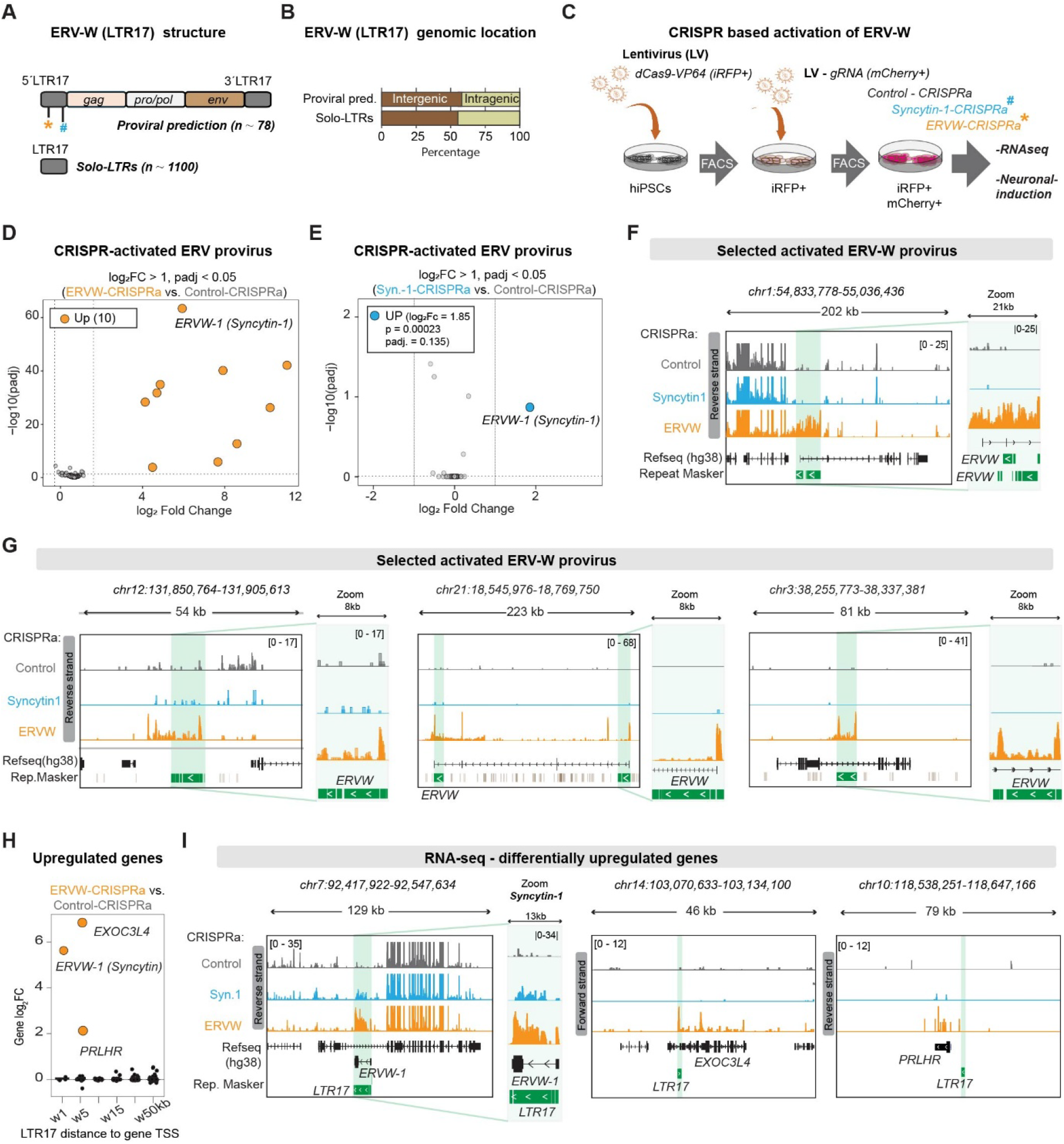
CRISPR mediated ERV-W activation in hiPSCs. (A) Structure of ERV-W elements with LTR17 (grey boxes) flanking the retroviral genes (*gag, pro, pol* and *env*). With proviral ERV-W predictions and solo-LTR17s estimations. * CRISPRa-ERVW and ^#^ CRISPRa-Syncytin-1 guides target location. (B) Genomic locations (inter-versus intragenic) of LTR17-ERV-Ws and solo-LTR17s based on the ERV RepeatMasker transcript annotation, 57.6 and 55.3% respectively. (C) Schematic of the CRISPR activation workflow and generation of three hiPSC-iNGN-cell lines: Control-CRISPRa, ERVW-CRISPRa and Syncytin-1-CRISPRa. Volcano plot of differential proviral ERV expression (DESeq2) following (D) ERV-W-CRISPRa and (E) Syncytin1-CRISPRa and summary of the differential expression analysis (DEA). Ten significantly upregulated ERV loci with seven being LTR17-ERV-Ws, including *ERVW-1* (Syncytin-1), no downregulated ERVs were detected (DESeq2 Wald test; |log₂FC| > 1, padj < 0.05). (F-G) Genome browser snapshots of selected CRISPR-activated ERV-W loci identified by bulk RNA-seq. Tracks show RPKM normalized reverse transcription for Control-CRISPRa (gray), Syncytin-1-CRISPRa (blue) and ERV-W-CRISPRa (orange). Green boxes indicate activated ERV-W loci (zoomed at right). (H) Expression changes (log₂FC) of genes located 1– 50 kb from the nearest LTR17 following ERV-W CRISPRa. (I) Genome browser tracks showing RPKM normalized expression of upregulated genes (*ERVW-1*, *EXOC3L4*, and *PRLHR*) together with the corresponding activated LTR17 element.

We designed CRISPR activation (CRISPRa) guide-RNAs (gRNA) to target the LTR17 consensus sequence (*AATGATCGGGATATAAACCC*, ERVW-CRISPRa). In addition, because *ERVW1* an LTR17-containing domesticated *env*-gene encoding the fusogenic protein Syncytin-1 could potentially contribute to downstream phenotypes, we designed a second gRNA (GCTAATTAGGCAAAGACAGG), termed Syncytin-1-CRISPRa, to selectively activate *ERVW1* as an additional control (Figure 2C). As a control we used a gRNA targeting LacZ (Control-CRISPRa), a sequence absent from the human genome. hiPSCs were first transduced with a lentiviral construct encoding dCas9-VP64 fused to an infrared reporter (iRFP) and subsequently isolated by fluorescence-activated cell sorting (FACS). In a second step, the sorted cells were transduced with a lentiviral vector encoding the gRNA, as well as a mCherry reporter, after which double-positive cells were obtained by FACS (Figure 2C). CRISPRa hiPSCs displayed characteristic colony morphology and positive OCT4 staining and reserved common cellular markers of pluripotency (Figure S2A, B), demonstrating that CRISPRa engineering did not alter cellular identity prior to differentiation.

RNA-seq analysis of ERVW-CRISPRa transduced hiPSCs revealed robust upregulation of seven ERV-W proviruses (Figure 2D). In contrast, Syncytin1-CRISPRa hiPSCs showed specific upregulation of *ERVW-1,* with no detectable upregulation of other ERV loci (Figure 2E). In control-CRISPRa hiPSCs, ERV-W expression was almost completely absent. Six of the seven upregulated ERV-W loci in ERVW-CRISPRa hiPSCs were intragenic, with three positioned in sense orientation relative to their host genes, including *ERVW1* (Figure 2E-G, I). Furthermore, a global expression analysis of all TEs confirmed that the largest categories of upregulated loci using repeatMasker annotations upon ERVW-CRISPRa were either LTR17 or HERVW-int (Figure S2C), confirming the specificity of our approach.

We next asked whether any of the upregulated ERV-W proviruses retained protein-coding potential. To address this, we examined the integrity of open reading frames (ORFs) within the activated ERV-W loci by assessing the presence of frameshift mutations and premature stop codons and identified five such regions containing at least one intact ORF, predominantly *pro*, only *ERVW1* (*HERV_dup2906*) retained an intact *env* (Table 1). These results indicate that five of the reactivated ERV-W loci retains the capacity to encode ERV-W derived peptides. Furthermore, since LTRs are known to act as gene enhancers and alternative promotors^27,36,37^ we examined the potential *cis*-acting transcriptional effects of ERV-W activation in hiPSCs by performing differential gene expression analysis comparing ERVW-CRISPRa to Control-CRISPRa (Figure S2D, E). We found that 7 genes were significantly affected (3 up- and 4 downregulated; padj <0.05, log2 fold change >1) upon ERVW-CRISPRa. Notably, the upregulated genes *EXOC3L4* and *PRLHR* upon ERVW-CRISPRa were located within 10 kb of an LTR17 element, indicating *cis*-regulatory effects of ERV-W activation (Figure 2H,I). In contrast, since the Syncytin-1 locus *ERVW1* is an intact ERV-W provirus that retains its LTRs and Env open reading frame, it is transcriptionally driven by its own LTR as expected.

**Table 1.** Open reading frames among CRISPR-activated ERV-Ws in hiPSCs. Sequence extracted with bedtools → All analysis steps were executed using SweepDNA followed by SweepScripts to process LTRID, ORFID and RetroVID^41^

| *Chr* | *Start* | *End* | *ID* | *Strand* | *gag* | *pro* | *pol* | *env* |
| --- | --- | --- | --- | --- | --- | --- | --- | --- |
| chr1 | 54906345 | 54914899 | *HERV_dup943*<br>(*ERV-W*) | - |  |  | 14 shifts,<br>8 stops | 3 shifts, 4<br>stops |
| chr1 | 54906754 | 54917807 | *HERV_dup944*<br>(*ERV-W*) | - | 2 shifts, 6<br>stops | **0 shifts, 0**<br>**stops** | 11 shifts,<br>2 stops | 3 shifts, 4<br>stops |
| chr3 | 38290491 | 38294225 | *HERV_dup1916*<br>(*ERV-W*) | - |  | **0 shifts, 0**<br>**stops** | 2 shifts, 2<br>stops |  |
| chr7 | 92466864 | 92478137 | *HERV_dup2906*<br>(*ERV-W*) | - |  | 2 shifts, 0<br>stops | 7 shifts, 4<br>stops | **0 shifts, 0**<br>**stops** |
| chr12 | 131867287 | 131880952 | *HERV_dup515*<br>(*ERV-W*) | - | 2 shifts, 5<br>stops | **0 shifts, 0**<br>**stops** | 10 shifts,<br>13 stops |  |
| chr12 | 58849353 | 58860023 | *HERV_dup637*<br>(*ERV-W*) | - | 4 shifts, 7<br>stops | **0 shifts, 0**<br>**stops** | 4 shifts, 7<br>stops |  |
| chr21 | 18753834 | 18760570 | *HERV_dup1435*<br>(*ERV-W*) | - | 5 shifts, 4<br>stops | 2 shifts, 1<br>stop | 3 shifts,<br>12 stops |  |

Taken together, these findings demonstrate robust and selective activation of multiple ERV-W loci in human iPSCs while maintaining cellular identity. The inclusion of a Syncytin-1-specific control enables discrimination between the effects of broader ERV-W activation and activation of the domesticated *ERVW-1* locus, making this a potent experimental set up for investigating the functional consequences of ERV-W expression in human neurons.

### ERV-W activation preserves neuronal identity but reduces spontaneous neuronal activity

We next used our forward neuronal reprogramming strategy in combination with CRISPRa to generate excitatory neurons overexpressing ERV-W and sampled cells for bulkRNA, ICC and electrophysiology at 4, 30 and ∼60 dpi (Figure 3A). The expression of stage-specific cell identity markers throughout differentiation from iPSCs to iNGNs was not affected by ERVW-CRISPRa and followed the expected developmental trajectory when compared to the two control conditions (Control-CRISPRa and Syncytin1-CRISPRa, Figure S3A). Pluripotency markers such as *NANOG* and *POU5F1* were rapidly downregulated upon iNGN induction, whereas neuronal markers including *DCX* and *MAP2* increased as cells acquired a post-mitotic neuronal identity already at 4dpi. We detected 95–98% vGlut1-positive excitatory neurons at 60 dpi by ICC of which all expressed mCherry, confirming stable gRNA expression throughout neuronal differentiation (Figure S3B). CRISPRa-modified neuronal cultures exhibited characteristic neuronal morphology with well-defined soma, widespread neurite outgrowth, and extensive network formation as shown by MAP2 staining at 60dpi (Figure 3B). Consistent with this, cleaved Caspase-3 immunoreactivity remained low and comparable between conditions at 75 dpi (Figure S3C), together indicating no detectable differences in cell viability. Furthermore, bulk RNA-seq analysis from six independent iNGN inductions (Control-vs. ERVW-CRISPRa) revealed reproducible ERV-W activation following differentiation into glutamatergic neurons, further validating sustained CRISPRa-mediated transgene expression (Figure S3D, E).

**Figure 3.**
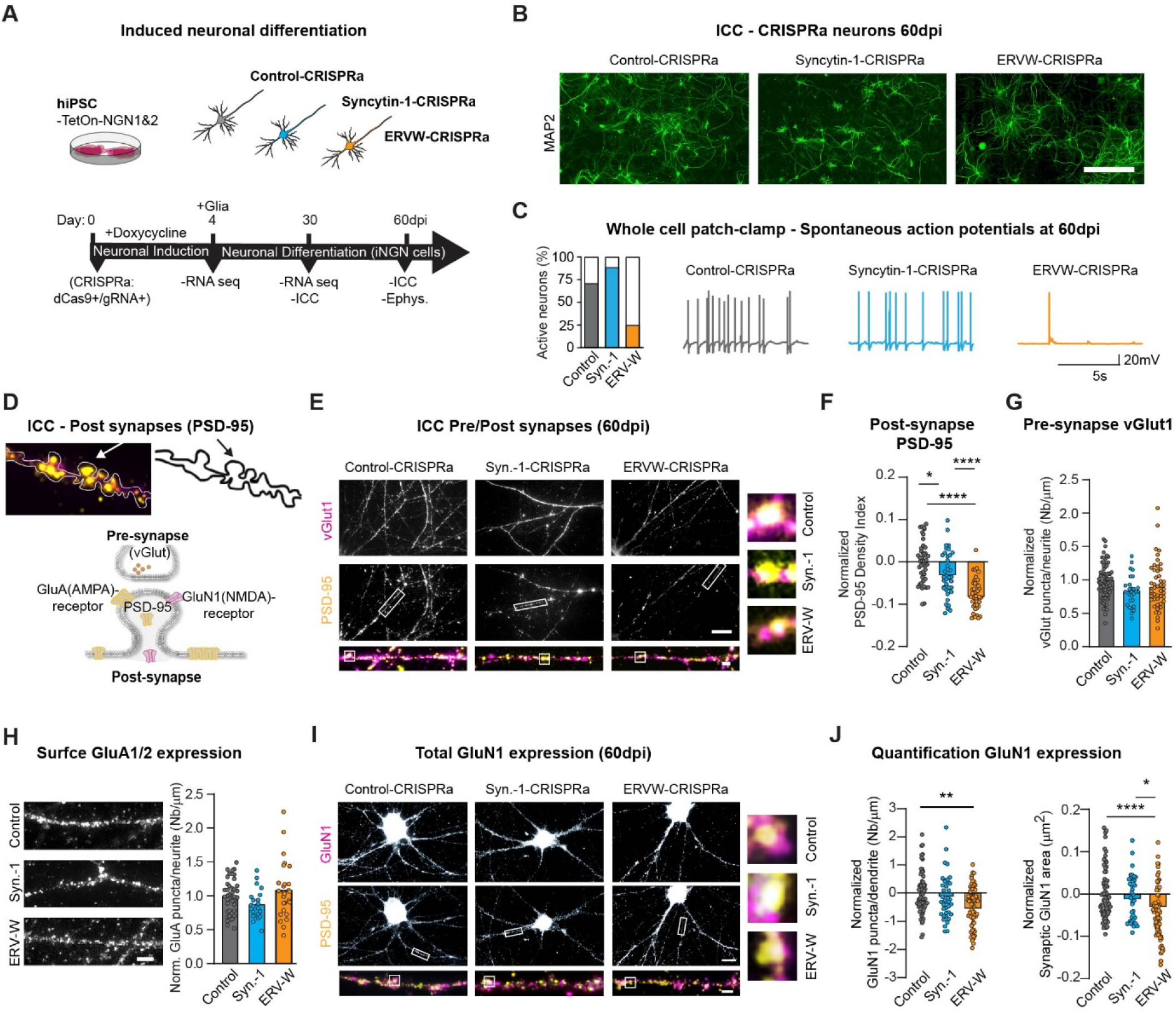
ERV-W activation disrupts excitatory synapse organization. (A) Schematic of neuronal induction and sample collection from CRISPR-modified hiPSCs. (B) Representative MAP2 immunostaining of neuronal cultures at 60 dpi. Scale bar, 50 μm. (C-left) % active neurons. Control-CRISPRa (9 neurons out of 13), Syncytin-1-CRISPRa (7/8) and ERV-W-CRISPRa (3/12) (C-right) Representative cell-attached recordings of spontaneous action potential currents from Control-CRISPRa (gray), Syncytin-1-CRISPRa (blue) and ERV-W-CRISPRa (orange) neurons. (D) Representative PSD-95-stained dendrite with spine contours at 60 dpi (top). Schematic of key synaptic proteins (bottom). (E) Representative vGlut1 (magenta) and PSD-95 (yellow) immunostaining. Colocalized puncta (white) indicates stable excitatory synapses. Insets show enlarged dendritic regions and individual synapses. Scale bars, 20 μm. (F) ERV-W CRISPRa reduced the postsynaptic density index ((PSD-95 puncta × area)/dendrite), (n = 39–45 dendrites from 20 cells/condition and 2 independent inductions, 56–72 dpi) ****p < 0.0001, *p = 0.011. (G) Presynaptic vGlut1 puncta density was unchanged following ERV-W CRISPRa. Dots represent puncta/neurite (Control-CRISPRa, n = 71 dendrites/50 cells/5 independent inductions, Syncytin1-CRISPRa n=29/20/2, ERV-W-CRISPRa n=46/30/3, 52–70 dpi), bars show means normalized to each induction. (H) Left: Representative surface GluA1/2 stainings and quantifications showing comparable AMPA receptor levels across conditions (n = 21-47 dendrites from 30 cells/condition and 3 independent inductions, 59–70 dpi). Scale bar, 5 μm. (I) Representative GluN1 (magenta) and PSD-95 (yellow) immunostaining. Colocalized puncta (white) indicate synaptic NMDA receptors, enlarged regions are shown below and at right. Scale bar, 20 μm. (J) ERV-W CRISPRa reduced both total GluN1 puncta/dendrite and synaptic GluN1 colocalization with PSD-95. Dots represent individual dendrites, bars show means normalized to induction (Control-CRISPRa, n = 68 dendrites /30 cells/3 independent inductions, Syncytin1-CRISPRa n=39/20/2, ERV-W-CRISPRa n=65/30/3, 56–72 dpi). ****p < 0.0001, **p = 0.0013, *p = 0.0185. Brown–Forsythe test/one-way ANOVA with Bonferroni correction.

To investigate the functional consequences of ERVW activation, we first evaluated whole-cell current-clamp recordings at 60dpi (Figure 3C and S4A). This revealed a marked reduction in spontaneous neuronal activity following ERV-W activation. Whereas 69 and 87% of control iNGNs (Control-CRISPRa and Syncytin1-CRISPRa) displayed spontaneous firing, only 25% of ERVW-CRISPRa iNGNs were spontaneously active (Figure 3C-left). Consistently, ERVW-CRISPRa iNGNs displayed fewer spontaneous action potentials, indicative of reduced neuronal excitability (Figure 3C-right). Because spontaneous firing reflects the integrated contribution of intrinsic membrane excitability and synaptic function, we next sought to determine whether ERV-W activation altered either property. Notably, ERVW-CRISPRa and control iNGNs showed comparable passive membrane properties with no differences in membrane capacitance nor input resistance between conditions and were morphologically similar as measured by automated segmentation of soma area (Figure S4B,C).

Next, we examined intrinsic membrane excitability to determine whether changes in neuronal firing could be attributed to alterations in the ability of individual neurons to generate action potentials. Whole-cell current-clamp recordings at 48–72 dpi revealed a more depolarized resting membrane potential in ERVW-CRISPRa iNGNs compared with both control conditions (Figure S4D). When held at -70 mV by constant current injection, ERVW-CRISPRa iNGNs still fired fewer action potential in response to depolarization current steps (Figure S4E,F), indicating that their reduced excitability is not secondary to the depolarized resting potential. Action potentials in ERVW-CRISPRa iNGNs were also smaller in peak amplitude, broader at half-width, and arose from a more depolarized threshold (Figure S4E,F), indicating changes in the cellular machinery that supports excitability. In voltage clamp, peak voltage-gated sodium currents were reduced (Figure S4G,H) whereas, voltage-gated potassium currents did not differ significantly between conditions at +30mV (Control-CRISPRa (mean = 3523, SEM = 516, n = 13), Syncytin-1-CRISPRa (mean = 4001, SEM = 658, n = 8) and, ERV-W-CRISPRa (mean = 2261, SEM = 249, n = 12).

Together, these findings reveal disruption of intrinsic neuronal excitability that may contribute to the reduced neuronal activity observed following ERV-W activation.

### ERV-W activation selectively disrupts postsynaptic organization in human excitatory neurons

Given the contribution of synaptic input to neuronal activity and the previous evidence of altered glutamate receptor function following exposure to an ERV-derived *env* protein^32^, we next examined the structural and functional organization of excitatory synapses. NMDA receptor signaling is a major determinant of synaptic plasticity yet has been reported to contribute only modestly to neuronal excitability in human iPSC-derived neurons^43,47^. Therefore, we first established whether a measurable NMDA receptor component could be detected in our human iNGN model system. Pharmacological blockade with D-APV (50μM) in control iNGNs at 60dpi produced a clear reduction in activity measured both by calcium imaging in Mg²⁺-free condition (Figure S5A, B) and by whole-cell patch-clamp recordings (Figure S5C), demonstrating a functional NMDA receptor contribution to neuronal signaling in our system. Together with previous reports^42,43,48^, these findings validate our human iNGNs as a suitable platform for investigating glutamatergic synaptic integrity.

We first investigated whether ERV-W activation altered the molecular organization of excitatory glutamatergic synapses. Specifically, we quantified vGlut1-positive puncta as a measurement of presynaptic glutamatergic connectivity and PSD-95 as a central component of the postsynaptic density involved in synaptic stabilization and receptor organization. Consistent with the characterization of the iNGNs (Figure S1), CRISPRa modified iNGNs exhibited a mature glutamatergic synaptic architecture, characterized by abundant PSD-95-positive postsynaptic structures, vGlut1-positive presynaptic terminals, and well-developed dendritic protrusions as shown at 60dpi (Figure 3B,D,E). We quantified PSD-95 staining from three independent inductions and found a significant reduction in the PSD-95 density index in ERV-W-CRISPRa iNGNs compared with Control-CRISPRa iNGNs, evident at 30 dpi (Control-CRISPRa: normalized mean = 1, SEM = 0.0077, n=37dendrites/30cells/3inductions; ERV-W-CRISPRa: normalized mean = -0.029, SEM = 0.0071, n=43/30/3; **p = 0.0072, Student’s *t*-test). This reduction persisted at 60 dpi, with ERV-W-CRISPRa iNGNs showing reduced PSD-95 density index compared with both control conditions (Figure 3E,F).

In contrast, quantitative analysis of immunofluorescence staining from three independent inductions revealed no significant change in the density of vGlut1-positive puncta along neurites following ERV-W activation (Figure 3E,G), indicating preserved presynaptic excitatory input. Together, these findings indicate that ERV-W activation is associated with a persistent reduction in postsynaptic PSD-95 organization, with no corresponding change in the density of presynaptic excitatory inputs, consistent with a selective alteration of postsynaptic organization and/or synaptic stability in ERV-W-expressing iNGNs.

Given that PSD-95 is a core organizer of the postsynaptic density that governs the nanoscale positioning and stabilization of glutamatergic receptors^6^, we next examined whether the observed structural alterations were accompanied by changes in synaptic receptor composition. While surface staining revealed comparable levels of AMPA receptors across all conditions (Figure 3H), total NMDA receptor levels were significantly reduced in ERVW-CRISPRa iNGNs compared to Control-CRISPRa and to both Control- and Syncytin1-CRISPRa in the synaptic compartment (Figure 3I, J). Notably, NMDA receptors are directly anchored to PSD-95 through interactions with the C-terminal tails of GluN2 subunits in contrast to AMPA receptors that has an indirect relation to PSD-95^5^. Because total staining captures both membrane-associated and intracellular receptor pools, these data do not resolve membrane accessible receptor localization but indicate an overall reduction in NMDA receptor abundance across cellular compartments. Together with the reduction in PSD-95, these findings point to a selective remodeling of postsynaptic composition in ERV-W expressing neurons, characterized by preserved membrane availability of AMPA receptors but reduced NMDA receptor expression and altered postsynaptic scaffold organization.

### ERV-W activation disrupts excitatory postsynaptic currents

Based on the observed reduction in PSD-95 and NMDA receptor expression, we next asked whether these molecular changes translated into altered postsynaptic responsiveness to glutamatergic stimulation. Because no differences were detected between ERV-W-CRISPRa and Syncytin-1-CRISPRa iNGNs in intrinsic electrophysiological or passive membrane properties, or in the postsynaptic phenotype assessed by immunocytochemistry, we focused subsequent synaptic electrophysiological analyses on the ERV-W-CRISPRa and Control-CRISPRa conditions. To first explore glutamatergic receptor function independently of presynaptic neurotransmitter release, focal glutamate was applied while iNGNs were voltage-clamped alternately at +40 and -70 mV (Figure S6A). Comparing responses at these voltages provides an approximate measure of the relative contribution of AMPA and NMDA receptors to the excitatory postsynaptic current, with lower values representing less NMDA receptor activity. Although not statistically conclusive, the mean +40/−70 mV AUC ratio was 0.5 in Control-CRISPRa iNGNs and 0.3 in ERVW-CRISPRa iNGNs (Figure S6B,C). Given that focal glutamate application activates both synaptic and extrasynaptic glutamate receptors, we next examined spontaneous excitatory postsynaptic currents (sEPSCs) to specifically assess glutamatergic receptor function at the synapse in ERV-W expressing iNGNs.

We first analyzed mixed AMPA- and NMDA-receptor mediated outward sEPSCs at +40 mV (Figure 4A). Event frequency was calculated from all detected events per cell, whereas amplitude and kinetic parameters were derived from isolated events to avoid contamination by overlapping synaptic activity. Neither the number of isolated events per cell (Figure 4B, p=0.29), nor the overall sEPSC frequency (Figure 4D, p=0.35) differed significantly between conditions. However, isolated events in ERVW-CRISPRa iNGNs displayed a larger amplitude compared with Control-CRISPRa iNGNs (Figure 4E-left), indicating stronger individual synaptic responses. Rise time was unaffected (Control-CRISPRa (median=1.42; Range=2.77; n=93) and ERV-W-CRISPRa (median=1.16; Range=4.0; n=76), p=0.12, whereas decay was markedly faster in ERVW-CRISPRa iNGNs (Figure 4C,E-right). Given that NMDA receptors contribute to the slow component of synaptic currents, the shorter decay is consistent with reduced NMDA receptor contribution, in line with decreased NMDA receptor expression detected by ICC.

**Figure. 4.**
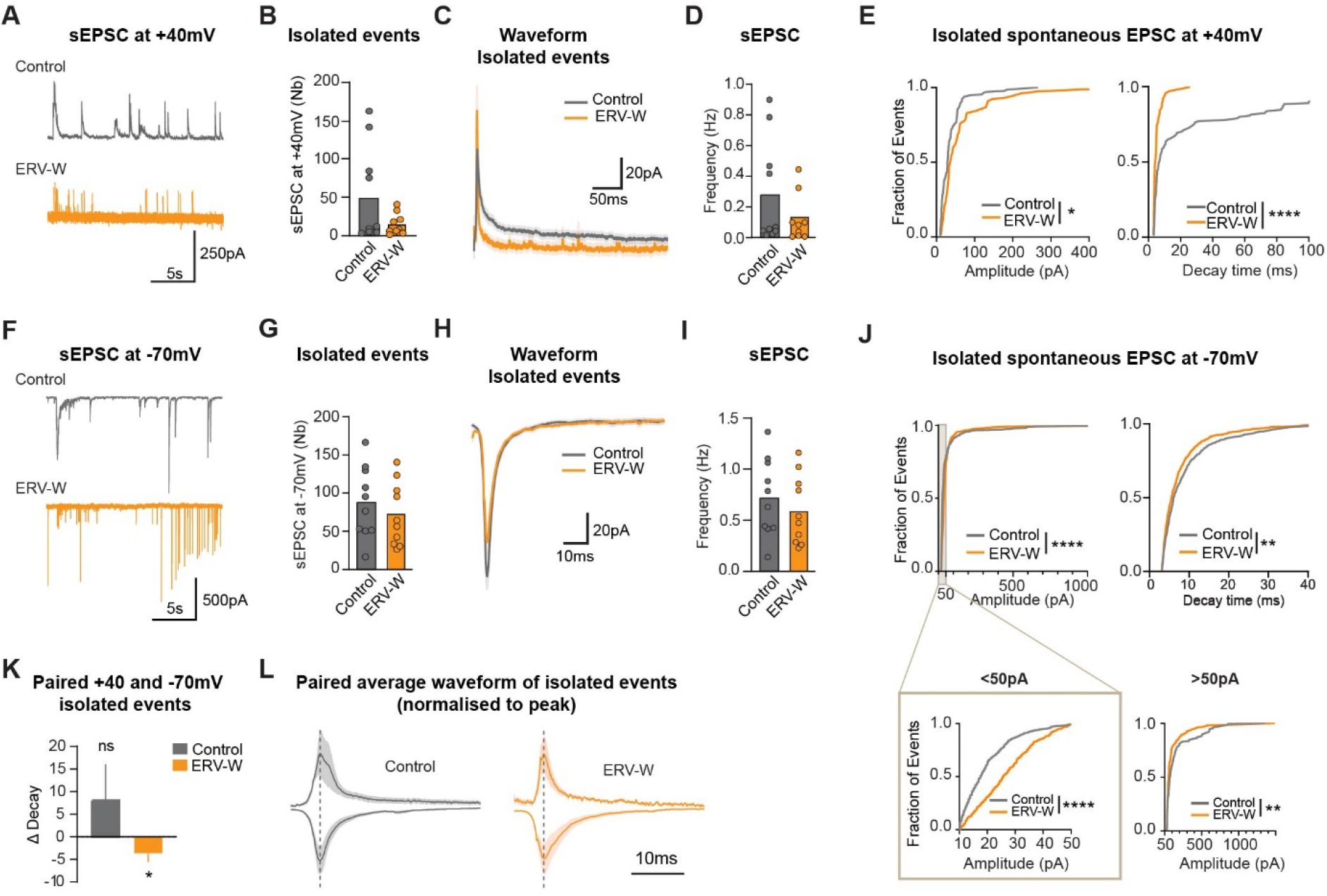
ERV-W activation perturbs post synaptic currents. (A) Representative traces of mixed AMPA- and NMDA receptor-mediated spontaneous excitatory postsynaptic currents (sEPSCs) recorded at +40 mV. (B) Number of isolated sEPSCs (isolation window ≥500 ms) per cell. (C) Mean sEPSC waveform ± SEM generated by cell-based averaging. (D) Mean sEPSC frequency per cell. (E) Cumulative distributions of isolated sEPSC amplitudes (left) and decay times (right). Events were pooled across cells (Control-CRISPRa, n = 10 neurons/131 events; ERV-W-CRISPRa, n = 9 neurons/76 events), **p = 0.01, ****p < 0.0001. (F) Representative traces of AMPA receptor-mediated sEPSCs recorded at −70 mV. (G) Number of isolated sEPSCs (isolation window ≥50 ms) per cell. (H) Mean sEPSC waveform ± SEM generated by cell-based averaging. (I) Mean sEPSC frequency per cell. (J) Cumulative distributions of isolated sEPSC amplitudes (left), ****p < 0.0001, including separate analyses of events <50 pA and >50 pA (bottom), **p = 0.0047 & ***p < 0.0001, and decay times (right),**p = 0.0033. Events were pooled across cells (Control-CRISPRa, n = 10 neurons/887 events; ERV-W-CRISPRa, n = 10 neurons/722 events). (K) Voltage-dependent change in decay time (Δdecay = decay₊₄₀ − decay₋₇₀) calculated from paired isolated events within individual neurons, *p = 0.047. (L) Mean paired waveforms of isolated events recorded at +40 and −70 mV, normalized to peak amplitude. Dots represent individual neurons; bars indicate mean ± SEM. Mann–Whitney, Kolmogorov–Smirnov, or Wilcoxon matched-pairs signed-rank tests, as indicated.

To examine AMPA receptor-mediated transmission in isolation, we next recorded inward currents at -70 mV (Figure 4F). The number of isolated events per cell did not differ between conditions (Figure 4G, p=0.41) nor did event frequency (Figure 4I, p=0.45). In contrast, the amplitude distributions of isolated events differed between ERVW-CRISPRa iNGNs compared to Control-CRISPRa iNGNs (Figure 4H,J-left). Among the events below 50 pA, which represents approximately 80% of all events in both groups, amplitudes were shifted towards larger values upon ERVW-CRISPRa, whereas higher-amplitude events (above 50pA) were shifted towards lower values (Figure 4J-bottom), indicating a redistribution of event amplitudes upon ERV-W activation. In addition, decay times were significantly shifted towards shorter values in ERVW-CRISPRa iNGNs compared to Control-CRISPRa (Figure 4J-right), indicating faster synaptic kinetics. These findings suggest that ERV-W expression does not detectably alter the frequency of glutamatergic receptor-mediated synaptic events, but instead alters their amplitude and decay kinetics, consistent with postsynaptic dysregulation and altered synaptic signaling dynamics.

Because decay kinetics differed at both holding potentials, we next asked whether this effect was also evident within individual neurons. We therefore used paired recordings, comparing responses at -70 mV and +40 mV within the same cell to minimize cell-to-cell variability. For each neuron, Δdecay was calculated from isolated events as the difference between the mean decay time at +40 mV and -70 mV (Δdecay = decay₊₄₀ − decay₋₇₀). Because NMDA receptor activation at depolarized potentials is expected to prolong synaptic decay, a positive Δdecay would be predicted. Control-CRISPRa iNGNs showed a positive Δdecay, consistent with this expected voltage-dependent NMDA receptor contribution, although the difference was not significantly different from zero (p=1.0). In contrast, ERVW-CRISPRa neurons showed a negative Δdecay, reflecting shorter decay at +40 mV than -70 mV within the same cells (Figure 4K,L). This reversal of the expected voltage dependence in ERVW-CRISPRa neurons suggests a reduced contribution of slow, NMDA receptor-mediated components to the synaptic currents.

Together, these findings support a model where ERV-W expression leaves excitatory synaptic connectivity largely intact while altering the organization of the postsynaptic compartment, with changes in PSD-95 and NMDA receptor abundance that are accompanied by altered synaptic strength and temporal integration.

## Discussion

ERVs comprise a substantial fraction of the human genome and have traditionally been considered largely transcriptionally silent in adult tissues, including the brain, where their expression is tightly restricted by epigenetic mechanisms such as DNA methylation. However, increasing evidence suggests that ERVs can become transcriptionally reactivated in neurological disease states and that specific cellular, inflammatory, and environmental stimuli may trigger their expression^15–17,30^. Despite these observations, the functional consequences of ERV activation in human neurons remain poorly understood.

Here, we provide direct evidence that transcriptional activation of ERV-W proviral sequences selectively disrupts postsynaptic organization and functional fine-tuning in maturing glutamatergic neurons. Although ERV-W expression has repeatedly been associated with a broad range of neurological and psychiatric disorders from studies in human samples, direct evidence for a causal contribution remains limited. Previous mechanistic studies have primarily focused on the potential effects of an ERV-W-derived envelope protein, using a full-length MSRV-type Env rather than a ERV-W gene product^32,40^. However, the genomic origin of this protein remains unresolved, with an ERV-W locus at Xq22.3 proposed as a candidate source^49,50^. Notably, ERVW-CRISPRa did not induce transcription of this locus (Figure S2F), indicating that the neuronal phenotypes observed here are unlikely to result from the activation of this particular locus. Rather, our findings point to several ERV-W loci and/or their transcriptional products as potential mediators of the observed effects on glutamatergic-synapse integrity. Despite targeting the consensus LTR17 sequence, only a small number of ERV-W elements were activated by our CRISPRa approach, indicating that local chromatin architecture, genomic context, and nearby regulatory elements influence the potential for different ERV-W loci to be expressed^17,34,36,51,52^.

We found that several of the CRISPRa-activated ERV-W loci retained intact open reading frames, raising the possibility that at least some of the observed phenotypes are mediated by ERV-W-derived peptides, increasingly recognized as potential contributors to neurological disease^15,31,38^. Emerging evidence suggests that ERV-derived peptides can engage immune pathways and serve as antigens for immune recognition. Aberrant expression of ERV-derived proteins has also been linked to protein aggregation and impaired synaptic function, providing potential mechanistic parallels to the neuronal phenotypes observed here^32,53,54^. However, establishing whether individual ERV-W loci give rise to functional protein products remains challenging. The development of robust reagents and approaches capable of detecting ERV-derived peptides with locus-level specificity will therefore be essential for determining whether, and through which mechanisms, disease-associated ERVs contribute to pathological processes.

Our results demonstrate that ERV-W activation results in molecular, structural, and electrophysiological changes in human neurons consistent with preferential disruption of postsynaptic organization and neuronal excitability. Rather than simply eliminating excitatory synapses, ERV-W activation appears to alter their molecular and functional properties. The co-occurrence of reduced PSD-95 abundance with altered NMDAR expression and glutamate-evoked responses is consistent with disrupted PSD-95-dependent organization of the postsynaptic compartment. PSD-95-containing scaffold complexes contribute to the nanoscale architecture of excitatory synapses and stabilize NMDARs at the postsynaptic membrane, thereby supporting their localization and function. Disruption of this organization affects NMDAR membrane retention and trafficking, with consequences for receptor localization and synaptic function without necessarily influencing synapse number^55–57^, suggesting that ERV-W activation may affect activity-dependent regulation of synaptic strength^58^. Further defining whether the reduction in PSD-95 density reflects reduced protein abundance at individual synapses, or changes in synapse number will provide important mechanistic insight into how ERV-W activation affects synaptic integrity.

At the functional level we identify two distinct consequences of ERV-W activation that converge on reduced neuronal activity, impaired intrinsic excitability and altered postsynaptic integration. Although these phenotypes may interact, it remains unclear whether they share a common mechanism or arise independently. The reduced intrinsic excitability indicates that decreased neuronal activity cannot be explained solely by altered postsynaptic integration, while changes in intrinsic excitability may in turn influence synaptic function. At the post synapse, altered glutamate-evoked responses are consistent with the reduced NMDAR abundance observed by ICC, suggesting that changes in NMDAR-mediated signaling may contribute to altered synaptic integration. The increased sEPSC amplitude despite preserved presynaptic measures may, in contrast, reflect a compensatory postsynaptic adaptation to reduced neuronal activity, consistent with homeostatic mechanisms that stabilize excitatory drive^59^. Importantly, unaffected presynaptic marker abundance and spontaneous event frequency does not by itself establish preservation of synapse number, structural connectivity, or presynaptic function. Thus, direct assessment of synapse number and pre–postsynaptic apposition, together with measurements of evoked synaptic transmission and presynaptic release probability, would provide a comprehensive assessment of structural and functional connectivity upon ERV-W activation. Such approaches would help distinguish selective postsynaptic remodeling from more subtle alterations in synaptic connectivity or presynaptic function.

In summary, ERV-W expression in human neurons reduces excitability and alters the efficiency with which existing excitatory synapses transmit and integrate information through disruption of a fundamental layer of postsynaptic organization. Given that impaired postsynaptic organization is recognized as a convergent feature across neurological and psychiatric disorders^5,9,60,61^ and that expression of ERVs are increasingly found in such disorders, our findings mechanistically link ERV-W activation and disease-relevant synaptic dysfunction. Our data suggest that diverse ERV-W loci and distinct modes of ERV-W activation may converge on similar neuronal outcomes despite substantial differences in their genomic origin. Together, these results provide a mechanistic framework for linking ERV-W activity to neuronal dysfunction that offers a potential causal explanation for its association with brain disease.

## Material and Methods

### Cell culture of human induced pluripotent stem cells

The commercially available human induced pluripotent stem cells (hiPSC) cell line (HPSI0114i-Kolf_2iei-C1; male; hereafter referred to KOLF2.1) was used in this study. Before experimentation, cells were karyotyped and immunoassayed as quality control measures. The hiPSCs were maintained on Matrigel matrix (Corning) diluted in DMEM/F12 (GIBCO) and 1% penicillin-streptomycin (P/S) (GIBCO) and plated on 6-well multidishes (Nunc [Thermo Fisher Scientific]) at 10000 cells/cm2 in mTeSRPlus supplemented media (STEMCELL Technologies) and 0.5% P/S. Cells were passaged every 3-4 days at 70-90% confluency. To passage, cells were rinsed once with Dulbecco’s Phosphate-Buffered Saline (DPBS) (GIBCO) and dissociated with Accutase (500 ul/well for 6-well plates, [GIBCO]) for 5 – 7 mins at 37°C. Dissociated cells were transferred to wash media (9.5 ml DMEM/F12, 500 µl Knockout Serum Replacement [GIBCO]) and centrifuged for 5 minutes at 400 x g. The cells were resuspended in mTeSRPlus media and plated with 10 µM Y276327 (Rock inhibitor (RI) [Miltenyi]) for expansion. Media was changed daily excluding RI. If applied, cells were frozen in CryoStore CS10 (STEMCELL technologies) media in cryotubes (Sarstedt) after disassociation as above. Cells were placed at -80°C for 48-72 hours in CoolCell boxes (Corning) before being placed at -150°C.

### iNGN-cells, neuronal induction

The human stem cell line KOLF2.1 was genetically modified by nucleofection with a plasmid containing a doxycycline-sensitive TetOn-Neurogenin 1 and 2 cassette for induced neuronal differentiation. These cells are called iNGN-cells.

To differentiate iNGN-cells to neurons we followed previously published protocols with minor modifications^42,43,62^. Briefly, iNGN-cells were differentiated for 4 days in a Matrigel hESC-Qualified Matrix (Corning)-coated 6-well plate. For neuronal induction, iNGN-cells were treated with 0.5 µg/ml doxycycline per day (Sigma) for 4 consecutive days. Ara-C (5 µM; cytosine b-D-arabinofuranoside hydrochloride, Sigma) was applied at 3 dpi to remove undifferentiated cells before re-seeding. At 4 dpi, cells were disassociated with Accutase for 5 minutes at 37°C, centrifuged at 400 g for 5 min, and carefully resuspended into a single cell suspension in Complete BrainPhys (BrainPhys Neuronal media [STEMCELL Technologies] with N2-A (STEMCELL Technologies), SM1 (STEMCELL Technologies), 1% P/S, ascorbic acid (200 µM [Sigma-Aldrich]), BDNF (20ng/ml [R&D]), and GDNF (20ng/ml [R&D])) that was equilibrated overnight 37°C in an incubator.

For biochemistry, neurons were reseeded onto poly-D-lysine (0.05 mg/ml [GIBCO]) and laminin-521(20 µg/ml [Biolamina]) coated 6-well plates (Nunc [Thermo Fisher Scientific]) at around 26,000 cells/cm2 in Complete BrainPhys media. For imaging and electrophysiology, the neurons were plated onto a 60 mm petri dish (Thermo Fisher Scientific) with four poly-D-lysine and laminin-521 coated 18 mm glass coverslips (Marienfeld) at around 7,000 cells/cm2 in Complete BrainPhys media.

Commercially available rat primary cortical astrocytes (GIBCO) were used in this study for co-culture with iNGNs. The astrocytes were thawed, expanded, and passaged according to manufacture instructions and maintained in Astrocyte Growth Medium (Thermo Fisher). The day before reseeding with iNGN cells, the astrocyte media was changed to Complete BrainPhys media. For neurons in 6-well plates, rat astrocytes (at around 40% confluence) plated on 22 mm glass coverslips (Marienfeld), with paraffin dots for separation, were flipped onto neurons after 2 hours. For neurons plated on paraffin dotted coverslips for imaging and electrophysiology; after 2 hours, the slips were flipped (neurons face-down) onto a 60 mm petri dish of rat astrocytes (at around 40% confluence) with Complete BrainPhys media. Half of the media was changed every 3-4 days using equilibrated Complete BrainPhys media. Ara-C was added for 24h at 7 dpi if needed

### CRISPR activation (CRISPRa)

First, we designed different guide RNAs (gRNAs): one gRNA targeting the LTR17 consensus sequence to activate the ERV-Ws (ERV-W-CRISPRa), a second gRNA selectively targeting ERVW1 (Syncytin-1-CRISPRa) and a control gRNA targeting LacZ, a sequence absent from the human genome (Control-CRISPRa). The guide sequences were inserted separately into a lentiviral backbone pLV hU6-sgRNA-mCherry using annealed oligos (see Table S1) and the BsmBI cloning site. For generation of a dCas9-VP64 cell line the dCas9 activation construct, we generated the lentiviral backbone pLV hUbC-VP64-dCas9-T2A-iRFP. The extended lentiviral production protocol has previously been published ^29,63^.

CRISPRa was established through two sequential lentiviral transductions. First, iNGN hiPSCs were transduced with lentiviral particles expressing VP64-dCas9-iRFP, followed by a second transduction with lentiviral particles expressing the gRNA-mCherry construct. To perform these transductions, cryopreserved iNGN cells (KOLF2.1 iNGN cell line, P8–10) were thawed as described above and seeded onto Matrigel-coated 12-well plates at a density of 71,500 cells/cm² in mTeSRPlus medium supplemented with RI. Medium was changed daily. Three days after seeding, cells were passaged and transduced with the first lentiviral construct encoding VP64-dCas9-iRFP in Matrigel-coated 12-well plates using mTeSR Plus medium supplemented with RI. Three days later, cells were passaged into Matrigel-coated 6-well plates in mTeSR Plus medium containing RI and 0.3 µg/mL puromycin. Medium supplemented with puromycin was replaced daily to eliminate cells lacking the iNGN construct. Four days after puromycin selection, iRFP-positive cells were isolated by fluorescence-activated cell sorting (FACSAria, BD Biosciences). Briefly, cells were detached and pelleted as described above, resuspended in mTeSR Plus medium supplemented with RI and Green Viability Dye (1:50,000; BD Via-Probe, BD Biosciences), and passed through a 70-µm cell strainer (Corning). Gating parameters were established using forward and side scatter to exclude debris and cell doublets. The iRFP-positive gates were set using untransduced hiPSCs. Sorting gates and strategies were validated via reanalysis of sorted cells with a requirement of > 97% purity cut-off. 100,000 iRFP+ and BD Via-Probe™ Green Nucleic Acid Stain negative cells were collected in mTeSR Plus media and plated for expansion. Once cells reached 70-90% confluency, they were either subjected directly to the second transduction or frozen.

Following the second transduction with the gRNA-mCherry constructs, VP64-dCas9-iRFP-positive cells were sorted by FACS to isolate double-positive (iRFP⁺/mCherry⁺/ViaGreen⁻) cells, which were subsequently expanded in culture. Once cultures reached 70–90% confluency, cells were cryopreserved. For all experiments, cells were thawed immediately before neuronal induction and were not expanded further. Guide efficiency was validated using bulk RNA sequencing, due to the repetitive nature of the targets (see bulkRNA analysis). A detailed protocol for lentiviral-mediated CRISPR modification of hiPSCs can be found at ^29,63^.

### Bulk RNA sequencing

Around 100 000 double-positive/BD Via-Probe™ Green-negative iNGN cells were collected for each sample, spun down at 1000 x rcf for 15 min at 10oC and snap-frozen on dry ice (for bulk RNA sequencing). For neuronal sampling, samples from different days post inductions (dpi) were collected from 3-6 different inductions. After removing the astrocyte cover glasses from the culture, and the neurons were briefly rinsed with DPDS (GIBCO), incubated with 350µl of RLT buffer (QIAGEN) with 1% β-Mercaptoethanol (Sigma-Aldrich) and mechanically scraped from the well for collection. Cell pellets or collected neurons were kept at -80oC until RNA was isolated. We isolated Total RNA using the RNeasy Mini Kit (QIAGEN) and the sequencing libraries were generated using Illumina TruSeq Stranded mRNA library prep kit (with poly-A selection) optimized for long library insert size and, sequenced on a Novaseq X plus (paired end 2 × 150bp).

### Bulk RNA sequencing analysis

Reads were uniquely mapped to hg38 reference genome using STAR aligner (version 2.7.8a; RRID:SCR_004463) using GENCODE version 38 as the guide GTF (--sjdbGTFfile, -- outFilterMultimapNmax 1, --outFilterMismatchNoverLmax 0.03). Alignment files were indexed (SAMtools version 1.16.1) and normalized bigwig files per strand were produced using bamCoverage (deeptools version 2.5.4) (--normalizeUsingRPKM --filterRNAstrand). Reads over genes, TEs (repeatMasker; open-4.0.5, filtered from tRNAs, simple repeats, small RNAs, and low-complexity regions), and retrotector predictions were quantified using featureCounts (subread package version 1.6.3; RRID:SCR_012919) forcing strandness (-s 2). Differential expression analyses were performed using DESeq2 (version 1.44.0; RRID: SCR_015687) with the corresponding features’ count matrices. DESeq2: lfcShrink was used to shrink fold changes. Gene size factors were calculated using median-of-ratios (DESeq2) and used as a normalization factor for TE and ERV expression. Expression data shown in heatmaps was normalized by gene size factors, summed to a pseudocount of 0.5, and log2-transformed.

### Immunocytochemistry

Live surface staining (10min at 37°C) was followed by 15 min fixation in 4% paraformaldehyde (PFA), quenching in 50mM NH4Cl for 10min, blocking for 30min in 1% bovine serum albumin (BSA) (Merck Millipore) and incubation with secondary antibodies coupled to Alexa fluorophores for 1h in 1% BSA at room temperature (R.T.). For staining of PFA fixed cells, incubation overnight with primary antibodies (see Table S1) at 4°C was done following an initial block and permeabilization step in 1% BSA and 0.05% Triton X-100 (Sigma-Aldrich) in phosphate-buffered saline (KPBS) for 1h. This was followed by washing and incubation with secondary antibodies as described above. The cells were finally washed and (if applicable) incubated with DAPI (1:1000 [Sigma-Aldrich]) in KPBS at room temperature for 10 minutes before being mounted with Mowiol (Calbiochem, Merck).

Images of neurons were randomly selected for analysis and collected on a motorized Nikon TiE total internal reflection microscopy (TIRF) with a Photometrix Prime95B sCMOS camera using a 100x objective and NIS-Elements software at the Lund Bioimaging Centre. Laser settings were maintained across conditions. For surface cluster analysis, dendritic branches were chosen manually (ROIs) and cluster areas and numbers were obtained using a manual threshold approach based on integrated fluorescence levels in ImageJ (NIH). Dendritic density of glutamatergic synapses (post-synaptic density index) was calculated as (number and area) of endogenous PSD-95 positive clusters/dendritic length and receptor abundance as cluster number/neurite length. For neuronal analysis, data was obtained from 2-4 independent neuronal inductions and later normalized within each induction. Imaging iPSCs and widefield images of neurons was conducted using a Leica epifluorescence microscope (DMI6000 B) and LasX software using 10x and 20x objectives. Imaging for soma size quantification was conducted using an Operetta CLS High-Content Analysis System (Revvity) followed by integrated analysis of specific settings of all nuclei and soma area settings across conditions. Synaptic spine images were acquired live on the TIRF microscope and using a Zeiss LSM 980 equipped with an Airyscan 2 (AX/AX R) detector (Carl Zeiss), operated in Airyscan mode to obtain enhanced-resolution confocal images of fixed PSD-95 neurons.

Figures were assembled in ImageJ (NIH) and Adobe Photoshop (Adobe Systems, San Jose, CA), only contrast and brightness were adjusted to optimize the image quality. All statistical analysis was performed in GraphPad Prism 11 (GraphPad Software, San Diego, USA). In brief, statistical significance was set at α = 0.05 after correction. Outliers were excluded using a ±2.5 SD criterion. Parametric statistical tests were applied when data passed the D’Agostino and Pearson’s omnibus normality test, hence two-tailed Student’s t test, or one-way ANOVA followed by Bonferroni’s multiple comparisons test were conducted. Statistical analysis can be found in detail in Table S2.

### Calcium imaging

For detailed calcium imaging protocol and analysis see Johansson et al. (2020)^32^. Briefly, 60dpi neurons transduced with GCaMP8f at 4dpi were transferred into Tyrode’s solution one day before imaging. The neurons were then moved to Mg^2+^-free Tyrode’s solution with 5 μM nifedipine (Tocris), 100μM Glutamate (Tocris) and 5 μM bicuculline (Tocris) 15 min before imaging. Time-lapse images were acquired with NIS-Elements software on a TiE total internal reflection microscopy (TIRF) with a Photometrix Prime95B sCMOS camera using a 100x objective. Two time-lapse movies were successively recorded at baseline and 5 min after bath application of 50 μM D-APV (Tocris).

### Patch-clamp electrophysiological recordings

Whole-cell patch-clamp electrophysiological recordings were performed at room temperature under continuous bath perfusion (2-3 mL/min) of an artificial cerebrospinal fluid (ACSF) containing (in mM): 119 NaCl, 2.5 KCl, 1.3 MgSO4, 2.5 CaCl2, 25 glucose, and 26 NaHCO3 continuously gassed with 95% O2 - 5% CO2, supplemented with 10 µM D-serine when spontaneous excitatory postsynaptic currents (sEPSC) were recorded.supplemented with 50 µM picrotoxin and 10 µM D-serine when spontaneous excitatory postsynaptic currents (sEPSC) were recorded. Picrotoxin was included to block GABA_A receptor-mediated currents, which would otherwise contribute large outward currents at +40 mV given the chloride reversal potential of approximately −70 mV imposed by the caesium-based internal solution. D-serine was included to saturate the glycine co-agonist binding site on NMDA receptors, ensuring maximal NMDA receptor availability during +40 mV recordings.

Human iPSC-derived neurons were selected based on mCherry fluorescence, viability and ability to obtain a stable seal and whole-cell configuration. Patch electrodes (filamented borosilicate capillary glass; 4 to 7 MΩ) were filled with an intracellular solution containing (in mM): 135.0 KMeSO4, 5.0 KCl, 0.5 CaCl2, 5.0 HEPES, 5.0 EGTA, 2.0 Mg-ATP, and 0.5 Na-GTP, adjusted to pH 7.30, osmolarity ∼290 mOsm. For the spontaneous EPSC recordings, the intracellular solution contained (in mM): 122.5 CsMeSO4, 8 NaCl, 10 HEPES, 0.2 EGTA, 2 MgATP, 0.3 Na3GTP; pH 7.3, osmolarity ∼290 mOsm. Caesium was used to block K+ channels and improve voltage-clamp quality. Liquid junction potentials were calculated using JPCalc (Clampex) and were approximately +7 mV for the K-methanesulfonate internal and +8 mV for the Cs-methanesulfonate internal against the ACSF bath. Junction potentials were not corrected, either online or offline. Nominal holding potentials of -70 mV and +40 mV thus corresponds to actual potentials of approximately -78 mV and +32 mV.

Immediately after break-in, resting membrane potential was recorded in current clamp (I = 0). Input resistance, membrane capacitance and access resistance were then obtained in voltage clamp from a 10mV membrane test from a holding potential of -70 mV; input resistance is reported as the steady-state resistance after subtraction of access resistance.

Current-Clamp recordings: Spontaneous firing was recorded for 3 min immediately after break-in in current clamp with no injected holding current. A cell was counted as spontaneously active if at least one action potential was fired during this recording period. To test intrinsic excitability, cells were held at –70 mV by constant current injection and action potentials were elicited with 250 ms depolarizing current steps from -50 to +350 pA in 10 pA increments. For each cell, the first action potential evoked at rheobase (the smallest current step eliciting a spike) was used to quantify AP properties. Threshold was defined as the membrane potential at which dV/dt first exceeded 20 mV/ms. Peak amplitude was measured from threshold to peak, and half-width as the action potential duration at half this amplitude.

Voltage-Clamp recordings: For voltage-gated current measurements, hiPSC-derived neurons were held at -70 mV in voltage clamp and 100-ms duration depolarizing steps were delivered starting from -70 to +40 mV in 10 mV increments. To minimize contamination by the capacitive transient, peak inward Na+ current was measured as the minimum current in a 2-50 ms window following step onset. Outward delayed rectifier K+ currents were measured as the maximum mean current in a 70-99 ms window after step onset. Spontaneous EPSC were recorded continuously for 3 minutes at each potential (−70 and +40 mV).

In some cells, agonist application was performed using a pressure ejection device (Picospritzer III, Parker). Puff pipettes were borosilicate electrodes (tip resistance 3-4 MΩ) filled with glutamate (100 µM) and D-serine (10 µM) dissolved in ACSF solution (adjusted to pH 7.30) and positioned under visual guidance at the soma or proximal dendrites at an approximate distance of 35 µm (range from 25 to 45 µm) from the cell membrane. 100-ms puffs were applied at 10 psi while holding cells at -70 mV in voltage clamp. An inter-puff interval of 15 s was used. Cells were classified as responsive if the peak current exceeded 3x baseline noise within a 500-ms window after puff onset. If no response was detected immediately, at least 5 puffs were delivered in ≥5 different locations. Once a clear response was identified, glutamatergic responses were recorded alternately at a holding potential of –70 mV and + 40 mV for a total of 5-10 puffs per potential. In a subset of cells (the last cell recorded on a given coverslip), to confirm NMDA receptor mediation of the puff-evoked current, D-APV (50 µM) was bath-applied after baseline puff.

Recordings were carried out with the Multiclamp 700B System and acquired with pClamp 11 (Molecular Devices). Signals were low-pass filtered at 2-5 kHz (Bessel, 4-pole) and digitized at 10 kHz for the spontaneous EPSC (10-20 kHz for other protocols). Series resistance was 11.4 ± 1.6, 15.5 ± 1.9 and 12.7 ± 2.1 MΩ in Control-, ERV-W- and Syncytin-1-CRISPRa neurons respectively (mean ± SEM; p=0.12), and was compensated by 70-80% (compensation bandwidth 5-10 kHz) and monitored throughout the recording. Recordings were rejected if Rs changed by >15%. No leak subtraction was used. Electrophysiology data were analyzed in Igor Pro (WaveMetrics) using the NeuroMatic package. All sEPSC analysis was performed using SynaptiQ v3, a custom software written in Python. The experimenter was blind to condition during recording and analysis (coverslips and files carried construct codes only, and the correspondence was not known by the experimenter during recording or analysis). Parametric statistical tests were applied when data passed the D’Agostino and Pearson’s omnibus normality test, hence two-tailed Student’s t test, or one-way ANOVA followed by Bonferroni’s multiple comparisons test were conducted. Comparisons between groups with data from non-Gaussian distributions were performed using the two-tailed unpaired Mann-Whitney U test.

### Spontaneous EPSC Detection and Analysis

Raw traces were baseline-corrected using a sliding-median algorithm (2-second window). This removes slow drift and holding current offset while preserving the fast transients of synaptic events. The corrected trace is used for all subsequent detection and measurement steps. Noise (σ_noise) was estimated from the baseline-corrected trace as the median absolute deviation of the sample-to-sample differences, scaled to an equivalent standard deviation (median 4.5 pA; not different between conditions, p ≥ 0.55). The threshold factor was determined for each dataset using a calibration procedure: a range of threshold factors (3-8×) was tested across all cells, and the factor minimizing the coefficient of variation (CV) of detected event counts across cells was selected as the most stable and reproducible setting. A threshold factor of 6× σ_noise was used for both -70 mV and +40 mV recordings in this study. Event onsets were detected where the smoothed first derivative of the low-pass-filtered trace (2 kHz) crossed a threshold scaled from σ noise. An event was retained if the signal at its peak exceeded 6 σ noise relative to the sliding-median baseline, and if its amplitude, measured from the local pre-event baseline to the peak, was at least 10 pA. Events were further required to meet the following criteria: Minimum 90-10% decay time: 3 ms; Dead time between events: 5 ms; post-event analysis window: 150 ms at -70 mV; 500 ms at +40 mV.

Threshold crossings with a 90–10% decay time below 3 ms were rejected outright and contribute to none of the reported measures, including event frequency. This criterion removed a substantial fraction of threshold crossings: 28% and 30% at −70 mV, and 41% and 59% at +40 mV (median across cells, Control-CRISPRa and ERVW-CRISPRa respectively; 26% and 44%, and 30% and 59%, of all threshold crossings pooled). The proportion rejected did not differ significantly between conditions (p = 0.85 at −70 mV, p = 0.20 at +40 mV). The rejected population had approximately symmetric kinetics, with a median rise/decay ratio near 1, whereas retained events showed the asymmetry expected of a synaptic conductance (median rise/decay ratio ∼ 0.2). Because this criterion truncates the decay distribution from below, the reported decay times describe the retained population only. To the extent that genuinely fast synaptic events fall below the 3 ms bound, decay times are overestimated and between-group differences attenuated. The longest measurable decay is set by the post-event analysis window.

The peak amplitude of each accepted event was characterized, and the decay was quantified as the 90–10% decay time: the interval between the timepoints at which the signal descends to 90% and 10% of peak amplitude, estimated by linear interpolation between adjacent samples. Events failing amplitude or decay criteria were rejected automatically; remaining events were available for manual review in the event gallery of the application.

Because synaptic events frequently occurred in bursts, and because decay time was long relative to typical inter-event intervals (particularly at +40 mV), a proportion of detected events overlapped with neighboring events. To obtain uncontaminated estimates of unitary synaptic properties, events were classified as isolated if no neighboring event occurred within a defined isolation window on either side: 150 ms at -70 mV and 500 ms at +40 mV. These windows were chosen to exceed the expected decay time at each voltage, ensuring that the baseline had fully recovered before and after the event of interest. sEPSC data used neuron as unit of analysis, and Mann–Whitney followed by Bonferroni-correction across six pre-specified measures per holding potential. For comparison of pooled events distributions, Kolmogorov-Smirnov test was used. Statistical analysis can be found in detail in Table S2.

## Acknowledgements

We thank Patrik Jern and Hervé Perron for comments and support and laboratory members for constructive discussions. We thank the strategic research areas MultiPark and StemTherapy for supporting the Joint Electrophysiology Core Facility and FACS facility at Lund University. We thank A. Hammarberg at the FACS facility and U. Jarl for technical assistance. We also thank the Bioimaging Centre (LBIC) and Center for Translational Genomics (CTG) at Lund University for providing expertise and service with imaging, sequencing and analysis.

## Funding

This work was supported by the Olle Engkvist foundation (E.J.), The Crafoord Foundation (E.J.), Forska utan djurförsök (E.J.), Åhlens-stiftelse (E.J.) y the Swedish Research Council (2022-02673 to J.J.), the Swedish Brain Foundation (FO2023-0232 to J.J.) and the Swedish Government Initiative for Strategic Research Areas (MultiPark & StemTherapy).

## Author contributions

Conceptualization: E.M.J. and J.J. Methodology: E.M.J. and J.J. Formal analysis/investigation: E.M.J., C.D-H., R.G., C.P. and J.Joh. Resources: E.M.J., J.J., and V.B. Writing (original draft): E.M.J. and J.J. Writing (review and editing): All authors. Supervision: E.M.J. and J.J. Project administration: E.M.J. and J.J. Funding acquisition: E.M.J. and J.J.

## Declaration of interests

The authors declare no competing interests.

## Declaration of generative AI and AI-assisted technologies in the manuscript preparation process

During the preparation of this work, the author(s) used GPT-5.6 Luna to check for grammatical incorrectness. The author(s) reviewed and edited the output as needed and take full responsibility for the content of the published article.

## Data and materials availability

All data needed to evaluate the conclusions in the paper are present in the paper and/or the Supplemental Information and the sequencing data presented in this study have been deposited to ENA accession number PRJEB128139. Original code and clean data for analyses is available at GitHub at https://github.com/Molecular-Neurogenetics/ERVW_Davis2026

## Notes

https://github.com/Molecular-Neurogenetics/ERVW_Davis2026

### Competing Interest Statement

The authors have declared no competing interest.

## References

1. Traynelis, S.F., Wollmuth, L.P., McBain, C.J., Menniti, F.S., Vance, K.M., Ogden, K.K., Hansen, K.B., Yuan, H., Myers, S.J., and Dingledine, R. (2010). Glutamate receptor ion channels: structure, regulation, and function. Pharmacol Rev 62, 405–496. 10.1124/pr.109.002451.

2. Vigneault, E., Poirel, O., Riad, M., Prud’homme, J., Dumas, S., Turecki, G., Fasano, C., Mechawar, N., and El Mestikawy, S. (2015). Distribution of vesicular glutamate transporters in the human brain. Front Neuroanat 9, 23. 10.3389/fnana.2015.00023.

3. Malenka, R.C., and Bear, M.F. (2004). LTP and LTD: an embarrassment of riches. Neuron 44, 5–21. 10.1016/j.neuron.2004.09.012.

4. Groc, L., and Choquet, D. (2020). Linking glutamate receptor movements and synapse function. Science 368. 10.1126/science.aay4631.

5. Paoletti, P., Bellone, C., and Zhou, Q. (2013). NMDA receptor subunit diversity: impact on receptor properties, synaptic plasticity and disease. Nat Rev Neurosci 14, 383–400. 10.1038/nrn3504.

6. Sheng, M., and Hoogenraad, C.C. (2007). The postsynaptic architecture of excitatory synapses: a more quantitative view. Annu Rev Biochem 76, 823–847. 10.1146/annurev.biochem.76.060805.160029.

7. Fromer, M., Pocklington, A.J., Kavanagh, D.H., Williams, H.J., Dwyer, S., Gormley, P., Georgieva, L., Rees, E., Palta, P., Ruderfer, D.M., et al. (2014). De novo mutations in schizophrenia implicate synaptic networks. Nature 506, 179–184. 10.1038/nature12929.

8. Selkoe, D.J. (2002). Alzheimer’s disease is a synaptic failure. Science 298, 789–791. 10.1126/science.1074069.

9. Südhof, T.C. (2018). Towards an Understanding of Synapse Formation. Neuron 100, 276–293. 10.1016/j.neuron.2018.09.040.

10. Taddei, R.N., and Duff, K.E. (2025). Synapse vulnerability and resilience across the clinical spectrum of dementias. Nat Rev Neurol 21, 353–369. 10.1038/s41582-025-01094-7.

11. Hoyt, S.J., Storer, J.M., Hartley, G.A., Grady, P.G.S., Gershman, A., de Lima, L.G., Limouse, C., Halabian, R., Wojenski, L., Rodriguez, M., et al. (2022). From telomere to telomere: The transcriptional and epigenetic state of human repeat elements. Science 376, eabk3112. 10.1126/science.abk3112.

12. Wells, J.N., and Feschotte, C. (2020). A Field Guide to Eukaryotic Transposable Elements. Annu Rev Genet 54, 539–561. 10.1146/annurev-genet-040620-022145.

13. Douville, R.N., and Nath, A. (2014). Human endogenous retroviruses and the nervous system. Handb Clin Neurol 123, 465–485. 10.1016/B978-0-444-53488-0.00022-5.

14. Duarte, R.R.R., Pain, O., Bendall, M.L., de Mulder Rougvie, M., Marston, J.L., Selvackadunco, S., Troakes, C., Leung, S.K., Bamford, R.A., Mill, J., et al. (2024). Integrating human endogenous retroviruses into transcriptome-wide association studies highlights novel risk factors for major psychiatric conditions. Nat Commun 15, 3803. 10.1038/s41467-024-48153-z.

15. Kury, P., Nath, A., Creange, A., Dolei, A., Marche, P., Gold, J., Giovannoni, G., Hartung, H.P., and Perron, H. (2018). Human Endogenous Retroviruses in Neurological Diseases. Trends Mol Med 24, 379–394. 10.1016/j.molmed.2018.02.007.

16. Garza, R., Sharma, Y., Atacho, D.A.M., Thiruvalluvan, A., Abu Hamdeh, S., Jonsson, M.E., Horvath, V., Adami, A., Ingelsson, M., Jern, P., et al. (2023). Single-cell transcriptomics of human traumatic brain injury reveals activation of endogenous retroviruses in oligodendroglia. Cell Rep 42, 113395. 10.1016/j.celrep.2023.113395.

17. Garza, R., Adami, A., Thiruvalluvan, A., Wijesinghe, S., Curle, A., Tam, O., Forcier, T., Lagka, D.A., Kazakou, N.L., Atacho, D.A.M., et al. (2026). Activation of transposable elements is linked to aregion- and cell type–specific interferon response in Parkinson’s disease. 10.1101/2025.09.03.673956.

18. Liu, X., Liu, Z., Wu, Z., Ren, J., Fan, Y., Sun, L., Cao, G., Niu, Y., Zhang, B., Ji, Q., et al. (2023). Resurrection of endogenous retroviruses during aging reinforces senescence. Cell 186, 287–304 e226. 10.1016/j.cell.2022.12.017.

19. Zhang, H., Li, J., Yu, Y., Ren, J., Liu, Q., Bao, Z., Sun, S., Liu, X., Ma, S., Liu, Z., et al. (2023). Nuclear lamina erosion-induced resurrection of endogenous retroviruses underlies neuronal aging. Cell Rep 42, 113396. 10.1016/j.celrep.2023.113396.

20. Steiner, J.P., Bachani, M., Malik, N., DeMarino, C., Li, W., Sampson, K., Lee, M.H., Kowalak, J., Bhaskar, M., Doucet-O’Hare, T., et al. (2022). Human Endogenous Retrovirus K Envelope in Spinal Fluid of Amyotrophic Lateral Sclerosis Is Toxic. Ann Neurol 92, 545–561. 10.1002/ana.26452.

21. Balestrieri, E., Arpino, C., Matteucci, C., Sorrentino, R., Pica, F., Alessandrelli, R., Coniglio, A., Curatolo, P., Rezza, G., Macciardi, F., et al. (2012). HERVs expression in Autism Spectrum Disorders. PLoS One 7, e48831. 10.1371/journal.pone.0048831.

22. Perron, H., Hamdani, N., Faucard, R., Lajnef, M., Jamain, S., Daban-Huard, C., Sarrazin, S., LeGuen, E., Houenou, J., Delavest, M., et al. (2012). Molecular characteristics of Human Endogenous Retrovirus type-W in schizophrenia and bipolar disorder. Transl Psychiatry 2, e201. 10.1038/tp.2012.125.

23. Guo, C., Jeong, H.H., Hsieh, Y.C., Klein, H.U., Bennett, D.A., De Jager, P.L., Liu, Z., and Shulman, J.M. (2018). Tau Activates Transposable Elements in Alzheimer’s Disease. Cell Rep 23, 2874–2880. 10.1016/j.celrep.2018.05.004.

24. Tam, O.H., Rozhkov, N.V., Shaw, R., Kim, D., Hubbard, I., Fennessey, S., Propp, N., Consortium, N.A., Fagegaltier, D., Harris, B.T., et al. (2019). Postmortem Cortex Samples Identify Distinct Molecular Subtypes of ALS: Retrotransposon Activation, Oxidative Stress, and Activated Glia. Cell Rep 29, 1164–1177 e1165. 10.1016/j.celrep.2019.09.066.

25. Li, W., Lee, M.H., Henderson, L., Tyagi, R., Bachani, M., Steiner, J., Campanac, E., Hoffman, D.A., von Geldern, G., Johnson, K., et al. (2015). Human endogenous retrovirus-K contributes to motor neuron disease. Sci Transl Med 7, 307ra153. 10.1126/scitranslmed.aac8201.

26. Buzdin, A., Kovalskaya-Alexandrova, E., Gogvadze, E., and Sverdlov, E. (2006). At least 50% of human-specific HERV-K (HML-2) long terminal repeats serve in vivo as active promoters for host nonrepetitive DNA transcription. J Virol 80, 10752–10762. 10.1128/JVI.00871-06.

27. Chuong, E.B., Elde, N.C., and Feschotte, C. (2016). Regulatory evolution of innate immunity through co-option of endogenous retroviruses. Science 351, 1083–1087. 10.1126/science.aad5497.

28. Faulkner, G.J., Kimura, Y., Daub, C.O., Wani, S., Plessy, C., Irvine, K.M., Schroder, K., Cloonan, N., Steptoe, A.L., Lassmann, T., et al. (2009). The regulated retrotransposon transcriptome of mammalian cells. Nat Genet 41, 563–571. 10.1038/ng.368.

29. Adami, A., Garza, R., Gerdes, P., Johansson, P.A., Dorazehi, F., Koutounidou, S., Castilla-Vallmanya, L., Atacho, D.A.M., Sharma, Y., Johansson, J.G., et al. (2025). LINE-1 retrotransposons mediate cis-acting transcriptional control in human pluripotent stem cells and regulate early brain development. Cell Genom 5, 100979. 10.1016/j.xgen.2025.100979.

30. Jönsson, M.E., Garza, R., Sharma, Y., Petri, R., Sodersten, E., Johansson, J.G., Johansson, P.A., Atacho, D.A., Pircs, K., Madsen, S., et al. (2021). Activation of endogenous retroviruses during brain development causes an inflammatory response. EMBO J 40, e106423. 10.15252/embj.2020106423.

31. Tamouza, R., Meyer, U., Foiselle, M., Richard, J.R., Wu, C.L., Boukouaci, W., Le Corvoisier, P., Barrau, C., Lucas, A., Perron, H., and Leboyer, M. (2021). Identification of inflammatory subgroups of schizophrenia and bipolar disorder patients with HERV-W ENV antigenemia by unsupervised cluster analysis. Transl Psychiatry 11, 377. 10.1038/s41398-021-01499-0.

32. Johansson, E.M., Bouchet, D., Tamouza, R., Ellul, P., Morr, A.S., Avignone, E., Germi, R., Leboyer, M., Perron, H., and Groc, L. (2020). Human endogenous retroviral protein triggers deficit in glutamate synapse maturation and behaviors associated with psychosis. Sci Adv 6, eabc0708. 10.1126/sciadv.abc0708.

33. Jern, P., and Coffin, J.M. (2008). Effects of retroviruses on host genome function. Annu Rev Genet 42, 709–732. 10.1146/annurev.genet.42.110807.091501.

34. Dopkins, N., O’Mara, M.M., Lawrence, E., Fei, T., Sandoval-Motta, S., Nixon, D.F., and Bendall, M.L. (2022). A field guide to endogenous retrovirus regulatory networks. Mol Cell 82, 3763–3768. 10.1016/j.molcel.2022.09.011.

35. Montserrat-Ayuso, T., Pujol, A., and Esteve-Codina, A. (2026). A comprehensive annotation of conserved protein domains in human endogenous retroviruses. NAR Genom Bioinform 8, lqag013. 10.1093/nargab/lqag013.

36. Brattås, P.L., Jönsson, M.E., Fasching, L., Nelander Wahlestedt, J., Shahsavani, M., Falk, R., Falk, A., Jern, P., Parmar, M., and Jakobsson, J. (2017). TRIM28 Controls a Gene Regulatory Network Based on Endogenous Retroviruses in Human Neural Progenitor Cells. Cell Rep 18, 1–11. 10.1016/j.celrep.2016.12.010.

37. Playfoot, C.J., Duc, J., Sheppard, S., Dind, S., Coudray, A., Planet, E., and Trono, D. (2021). Transposable elements and their KZFP controllers are drivers of transcriptional innovation in the developing human brain. Genome Res 31, 1531–1545. 10.1101/gr.275133.120.

38. Perron, H., Garson, J.A., Bedin, F., Beseme, F., Paranhos-Baccala, G., Komurian-Pradel, F., Mallet, F., Tuke, P.W., Voisset, C., Blond, J.L., et al. (1997). Molecular identification of a novel retrovirus repeatedly isolated from patients with multiple sclerosis. The Collaborative Research Group on Multiple Sclerosis. Proc Natl Acad Sci U S A 94, 7583–7588. 10.1073/pnas.94.14.7583.

39. Karlsson, H., Bachmann, S., Schroder, J., McArthur, J., Torrey, E.F., and Yolken, R.H. (2001). Retroviral RNA identified in the cerebrospinal fluids and brains of individuals with schizophrenia. Proc Natl Acad Sci U S A 98, 4634–4639. 10.1073/pnas.061021998.

40. Herrero, F., Heeb, C., Meier, M., Lin, H.Y., Mueller, F.S., Schalbetter, S.M., Gruchot, J., Weber-Stadlbauer, U., Notter, T., Perron, H., et al. (2025). Recapitulation and reversal of neuropsychiatric phenotypes in a mouse model of human endogenous retrovirus type W expression. Mol Psychiatry 30, 3325–3337. 10.1038/s41380-025-02955-9.

41. Sperber, G.O., Airola, T., Jern, P., and Blomberg, J. (2007). Automated recognition of retroviral sequences in genomic data -: RetroTector©. Nucleic Acids Research 35, 4964–4976. 10.1093/nar/gkm515.

42. Busskamp, V., Lewis, N.E., Guye, P., Ng, A.H., Shipman, S.L., Byrne, S.M., Sanjana, N.E., Murn, J., Li, Y., Li, S., et al. (2014). Rapid neurogenesis through transcriptional activation in human stem cells. Mol Syst Biol 10, 760. 10.15252/msb.20145508.

43. Lam, R.S., Topfer, F.M., Wood, P.G., Busskamp, V., and Bamberg, E. (2017). Functional Maturation of Human Stem Cell-Derived Neurons in Long-Term Cultures. PLoS One 12, e0169506. 10.1371/journal.pone.0169506.

44. Jern, P., Sperber, G.O., and Blomberg, J. (2004). Definition and variation of human endogenous retrovirus H. Virology 327, 93–110. 10.1016/j.virol.2004.06.023.

45. Gimenez-Orenga, K., and Oltra, E. (2021). Human Endogenous Retrovirus as Therapeutic Targets in Neurologic Disease. Pharmaceuticals (Basel) 14. 10.3390/ph14060495.

46. Sperber, G.O., Airola, T., Jern, P., and Blomberg, J. (2007). Automated recognition of retroviral sequences in genomic data--RetroTector. Nucleic Acids Res 35, 4964–4976. 10.1093/nar/gkm515.

47. Zhang, Y., Pak, C., Han, Y., Ahlenius, H., Zhang, Z., Chanda, S., Marro, S., Patzke, C., Acuna, C., Covy, J., et al. (2013). Rapid single-step induction of functional neurons from human pluripotent stem cells. Neuron 78, 785–798. 10.1016/j.neuron.2013.05.029.

48. Gameiro-Ros, I., Tengolics, A.J., Prytkova, I., Kamarajan, C., Pang, Z.P., Goate, A.M., Hart, R.P., and Slesinger, P.A. (2026). Uncovering NMDA receptor-mediated high firing activity in excitatory human neurons. iScience 29, 115061. 10.1016/j.isci.2026.115061.

49. Roebke, C., Wahl, S., Laufer, G., Stadelmann, C., Sauter, M., Mueller-Lantzsch, N., Mayer, J., and Ruprecht, K. (2010). An N-terminally truncated envelope protein encoded by a human endogenous retrovirus W locus on chromosome Xq22.3. Retrovirology 7, 69. 10.1186/1742-4690-7-69.

50. Charvet, B., Pierquin, J., Brunel, J., Gorter, R., Quetard, C., Horvat, B., Amor, S., Portoukalian, J., and Perron, H. (2021). Human Endogenous Retrovirus Type W Envelope from Multiple Sclerosis Demyelinating Lesions Shows Unique Solubility and Antigenic Characteristics. Virol Sin 36, 1006–1026. 10.1007/s12250-021-00372-0.

51. Ho, J.S.Y., Douse, C.H., and Marazzi, I. (2026). Transposable element DNA and RNA: Drivers of gene expression, evolution, and disease. Cell 189, 3513–3540. 10.1016/j.cell.2026.05.003.

52. Castro-Diaz, N., Ecco, G., Coluccio, A., Kapopoulou, A., Yazdanpanah, B., Friedli, M., Duc, J., Jang, S.M., Turelli, P., and Trono, D. (2014). Evolutionally dynamic L1 regulation in embryonic stem cells. Genes Dev 28, 1397–1409. 10.1101/gad.241661.114.

53. Liu, S., Heumuller, S.E., Hossinger, A., Muller, S.A., Buravlova, O., Lichtenthaler, S.F., Denner, P., and Vorberg, I.M. (2023). Reactivated endogenous retroviruses promote protein aggregate spreading. Nat Commun 14, 5034. 10.1038/s41467-023-40632-z.

54. Padmanabhan Nair, V., Liu, H., Ciceri, G., Jungverdorben, J., Frishman, G., Tchieu, J., Cederquist, G.Y., Rothenaigner, I., Schorpp, K., Klepper, L., et al. (2021). Activation of HERV-K(HML-2) disrupts cortical patterning and neuronal differentiation by increasing NTRK3. Cell Stem Cell 28, 1566–1581 e1568. 10.1016/j.stem.2021.04.009.

55. Bard, L., Sainlos, M., Bouchet, D., Cousins, S., Mikasova, L., Breillat, C., Stephenson, F.A., Imperiali, B., Choquet, D., and Groc, L. (2010). Dynamic and specific interaction between synaptic NR2-NMDA receptor and PDZ proteins. Proc Natl Acad Sci U S A 107, 19561–19566. 10.1073/pnas.1002690107.

56. Heine, M., Groc, L., Frischknecht, R., Beique, J.C., Lounis, B., Rumbaugh, G., Huganir, R.L., Cognet, L., and Choquet, D. (2008). Surface mobility of postsynaptic AMPARs tunes synaptic transmission. Science 320, 201–205. 10.1126/science.1152089.

57. Lin, Y., Skeberdis, V.A., Francesconi, A., Bennett, M.V., and Zukin, R.S. (2004). Postsynaptic density protein-95 regulates NMDA channel gating and surface expression. J Neurosci 24, 10138–10148. 10.1523/JNEUROSCI.3159-04.2004.

58. Carroll, R.C., and Zukin, R.S. (2002). NMDA-receptor trafficking and targeting: implications for synaptic transmission and plasticity. Trends Neurosci 25, 571–577. 10.1016/s0166-2236(02)02272-5.

59. Turrigiano, G.G., Leslie, K.R., Desai, N.S., Rutherford, L.C., and Nelson, S.B. (1998). Activity-dependent scaling of quantal amplitude in neocortical neurons. Nature 391, 892–896. 10.1038/36103.

60. Dupuis, J.P., Nicole, O., and Groc, L. (2023). NMDA receptor functions in health and disease: Old actor, new dimensions. Neuron 111, 2312–2328. 10.1016/j.neuron.2023.05.002.

61. Grant, S.G. (2012). Synaptopathies: diseases of the synaptome. Curr Opin Neurobiol 22, 522–529. 10.1016/j.conb.2012.02.002.

62. Habibey, R., Striebel, J., Sharma, K., and Busskamp, V. (2022). Optogenetic Control of Human Stem Cell-Derived Neurons. Methods Mol Biol 2501, 339–360. 10.1007/978-1-0716-2329-9_17.

63. 10.17504/protocols.io.8epv5ro85g1b/v1.

