## Supplementary material for "Endogenous retrovirus activation disrupts postsynaptic organization in human glutamatergic neurons": Table S1

Supplemental Table 1. Products and resources

| REAGENT or RESOURCE | SOURCE | IDENTIFIER |
|---|---|---|
| **Antibodies** | | |
| mouse anti-OCT3/4 | Sant Cruz | RRID: AB_628051 |
| rabbit anti-NANOG | Abcam | RRID: AB_446437 |
| chicken anti-MAP2 | Abcam | RRID:AB_2138153 |
| mouse anti-PSD-95 | Invitrogen | RRID:AB_2092361 |
| rabbit anti-VGLUT1 | SYSY antibodies | Cat# 135 302 |
| rabbit anti-GFP | Molecular Probes | RRID:AB_221570 |
| rabbit anti-GluN1 | Alomone labs | RRID:AB_2040023 |
| mouse anti-Synaptophysin | Millipore | RRID:AB_94786 |
| rabbit anti-Cleaved Caspase-3 | Cell Signaling | RRID:AB_2341188 |
| mouse anti-GluA1+2 | SYSY antibodies | Cat# 182 411 |
| donkey anti-mouse Alexa Fluor 568 | Invitrogen | RRID:AB_11180865 |
| donkey anti-rabbit Alexa Fluor 488 | Invitrogen | RRID:AB_2535792 |
| donkey anti-rabbit Alexa Fluor 568 | Invitrogen | RRID:AB_2534017 |
| goat anti-chicken Alexa Fluor 488 | Invitrogen | RRID:AB_2534096 |
| **Chemicals, peptides, and recombinant proteins** | | |
| Rock-inhibitor, Y27632 | Miltenyi Biotec | Cat# 130-106-538 |
| Penicillin-Streptomycin | GIBCO (ThermoFisher) | Cat# 15140122 |
| Knockout replacement serum (KSR) | GIBCO(ThermoFisher) | Cat# 10828010 |
| Bicuculline | Tocris | Cat# 0131 |
| D-AP5 | Tocris | Cat# 0106 |
| Nifedipine | Tocris | Cat# 1075 |
| mTeSR Plus | STEMCELL | Cat# 100-0276 |
| Laminin-521 | Biolamina | Cat# LN521 |
| StemPro Accutase | GIBCO(ThermoFisher) | Cat# A1110501 |
| Matrigel | Corning | Cat# 354277 |
| DMEM/F-12 | GIBCO(ThermoFisher) | Cat# 31330-038 |
| DPBS | GIBCO(ThermoFisher) | Cat# 14190086 |
| Puromycin Dihydrochloride | ThermoFisher | Cat# A1113803 |
| Astrocyte growth media | ThermoFisher | Cat# A126130 |
| CryoStor CS10 | STEMCELL | Cat# 7952 |
| Cytosine β-D-arabinofuranoside hydrochloride | Sigma-Aldrich | Cat# C6645 |
| BrainPhys Neuronal Medium N2-A and SM1 kit | STEMCELL | Cat# 05793 |
| Ascorbic acid | Sigma-Aldrich | Cat# A92902 |
| Recombinant Human BDNF Protein | R&D Systems | Cat# 11166-BD |
| Recombinant Human GDNF Protein | R&D Systems | Cat# 212-GD |
| Poly-D-Lysine | GIBCO(ThermoFisher) | Cat# A3890401 |
| Paraffin | Sigma-Aldrich | Cat# P7793 |
| Polyethyleneimine | Polysciences | Cat# PN 23966 |
| Paraformaldehyde | Sigma-Aldrich | Cat# 158127 |
| Bovine serum albumin | Sigma-Aldrich | Cat# A9418 |
| Mowiol | Sigma-Aldrich | Cat# 81381 |
| Doxycycline hydrochloride | Sigma-Aldrich | Cat# D3447 |
| Green Viability Dye | BD Biosciences | Cat# 565802 |
| Buffer RLT | QIAGEN | Cat# 79216 |

| | | |
|---|---|---|
| DRAQ7 | BD Biosciences | Cat# 564904 |
| β-Mercaptoethanol | Sigma-Aldrich | Cat# M6250 |
| Critical commercial assays | | |
| RNeasy Mini Kit | QIAGEN | Cat# 74104 |
| TruSeq Stranded mRNA LP | Illumina | Cat# 20020594 |
| **Deposited data** | | |
| https://github.com/Molecular-Neurogenetics/ERVW_Davis2026 | | |
| ENA accession number PRJEB128139 | | |
| **Experimental models: Cell lines** | | |
| HPSI0114i-Kolf_2-C1, male | Wellcome Trust Sanger Institute | RRID:CVCL_9S58 |
| hiPSC iNGN | This paper | N/A |
| Rat primary cortical astrocytes | Thermo Fisher | Catalog # N7745100 |
| hiNGN_ERV-W_CRISPRa | This paper | N/A |
| hiNGN_Control_CRISPRa | This paper | N/A |
| hiNGN_Syncytin-1_CRISPRa | This paper | N/A |
| **Oligonucleotides:** | | |
| ERV-W-CRISPRa guide | forward | **CACCG**AATGATCGGGATATAAACCC |
| | reverse | **AAAC**GGGTTTATATCCCGATCATT**C** |
| Syncytin-CRISPRa guide | forward | **CACCG**GCTAATTAGGCAAAGACAGG |
| | reverse | **AAAC**CCTGTCTTTGCCTAATTAGC**C** |
| Non-targeting control (LacZ) guide | forward | **CACCG**TGCGAATACGCCCACGCGAT |
| | reverse | **AAAC**ATCGCGTGGGCGTATTCGCA**C** |
| **Recombinant DNA** | | |
| pLV hU6-sgRNA-mCherry | This paper | N/A |
| pLV hUbC-VP64-dCas9-T2A-iRFP | This paper | N/A |
| TetOn-Neurogenin 1 and 2 cassette | Volker Busskamp | N/A |
| GCamp8f | Henrik Ahlenius | N/A |
| **Software and algorithms** | | |
| IgV (version 2.19.4) | http://www.broadinstitute.org/igv/ | RRID:SCR_011793 |
| ImageJ (version 2.16.0/1.54p) | NIH | RRID:SCR_003070 |
| NeuroMatic package is v3.0s | SciCrunch Registry | RRID:SCR_004186 |
| JPCalcW supplied with pClamp 11.3 | Molecular Devices | RRID:SCR_011323 |
| Igor Pro (version 9.05) | WaveMetrics | RRID:SCR_000325 |
| Adobe Photoshop (2023) | Adobe | RRID:SCR_014199 |
| Adobe Illustrator (2023) | Adobe | RRID:SCR_010279 |
| STAR aligner (version 2.7.8a) | Dobin et al | RRID:SCR_004463 |
| SAMtools (version 1.16.1) | Li et al | RRID:SCR_002105 |
| Deeptools (version 2.5.4) | Ramirez et al | RRID:SCR_016366 |
| RepeatMasker (open-4.0.5) | Sperber el al | RRID:SCR_012954 |
| featureCounts (version 1.6.3) | Liao et al | RRID:SCR_012919 |
| DESeq2 (version 1.44.0) | Love et al | RRID:SCR_015687 |
| GraphPad Prism 11 | GraphPad Software | RRID:SCR_002798 |
| MATLAB R2024a | MathWorks Inc. | RRID:SCR_001622 |
