## Supplementary material for "Endogenous retrovirus activation disrupts postsynaptic organization in human glutamatergic neurons": Table S2

**Supplemental Table 2.** Statistical details

| Figure | Panel | Data | | | Control-CRISPRa | | | Syncytin-1-CRISPRa | | | ERV-W-CRISPRa | | | Statisitics | | | | |
|---|---|---|---|---|---|---|---|---|---|---|---|---|---|---|---|---|---|---|
| **Figure 3** | | | N (Unit) | Unit | Mean | SD | N | Mean | SD | N | Mean | SD | N | Test | | Value test | Padj-value | |
| | **F** | vGlut/neurite | Neurite-Cells | Nb | 1 | 0,2889 | 71 | 0,8462 | 0,2772 | 29 | 0,9285 | 0,3565 | 46 | Brown-Forsythe test | Cont.-ERVW | F=2.659 | 0,6742 | ns |
| | | Normalised per exp. | | | | | | | | | | | | One-way ANOVA, Bonferroni | Cont.-Syn. | | 0,0772 | ns |
| | | | | | | | | | | | | | | | ERVW-Syn. | | 0,7918 | ns |
| | **G** | PSD-95 density index | Dendrite-Cells | | 0 | 0,04852 | 45 | -0,03015 | 0,05251 | 39 | -0,07575 | 0,03671 | 42 | Brown-Forsythe test | Cont.-ERVW | F=29.34 | <0,0001 | **** |
| | | Normalised per exp. | | | | | | | | | | | | One-way ANOVA, Bonferroni | Cont.-Syn. | | 0,0105 | * |
| | | | | | | | | | | | | | | | ERVW-Syn. | | <0,0001 | **** |
| | **H** | GluA1-2/neurite | Dendrite-Cells | Nb | 1 | 0,2515 | 41 | 0,8771 | 0,22 | 21 | 1,081 | 0,4127 | 27 | Brown-Forsythe test | Cont.-ERVW | F=2,679 | 0,8483 | ns |
| | | Normalised per exp. | | | | | | | | | | | | One-way ANOVA, Bonferroni | Cont.-Syn. | | 0,4058 | ns |
| | | | | | | | | | | | | | | | ERVW-Syn. | | 0,0696 | ns |
| | **J (left)** | GluN1/dendrite | Dendrite-Cells | Nb | 0 | 0,7395 | 68 | -0,1192 | 0,7922 | 39 | -0,4574 | 0,7037 | 65 | Brown-Forsythe test | Cont.-ERVW | F=6.674 | 0,0013 | ** |
| | | Normalised per exp. | | | | | | | | | | | | One-way ANOVA, Bonferroni | Cont.-Syn. | | 0,7015 | ns |
| | | | | | | | | | | | | | | | ERVW-Syn. | | 0,0642 | ns |
| | **J (right)** | Synaptic GluN1 area | Dendrite-Cells | $um^2$ | 0 | 0,1488 | 68 | -0,04711 | 0,1176 | 39 | -0,1202 | 0,1202 | 65 | Brown-Forsythe test | Cont.-ERVW | F=13.96 | <0,0001 | **** |
| | | Normalised per exp. | | | | | | | | | | | | One-way ANOVA, Bonferroni | Cont.-Syn. | | 0,1794 | ns |
| | | | | | | | | | | | | | | | ERVW-Syn. | | 0,0185 | * |
| Figure | Panel | Data | | | Control-CRISPRa | | | Syncytin-1-CRISPRa | | | ERV-W-CRISPRa | | | Statisitics | | | | |
| **Figure 4** | | | | | Median | Range | N | | | | Median | Range | N | Test | | Value test | Padj-value | |
| | **B** | Nb of isol. events | Cell | Nb | 9.5 | 161 | 10 | | | | 6 | 40 | 10 | Mann-Whitney,Two-tailed | | 35,5 | 0,2876 | ns |
| | | | | Unit | Median | Range | N | | | | Median | Range | N | | | | | |
| | **D** | Frequency | Cell | Hz | 0,0555 | 0,891 | 10 | | | | 0,077 | 0,443 | 9 | Mann-Whitney,Two-tailed | | 33 | 0,3455 | ns |
| | **E** | Amplitude | Event | pA | 28,69 | 255,4 | 131 | | | | 36,16 | 1120 | 76 | Kolmogorov-Smirnov | | 0,2346 | 0,01 | * |
| | | Rise Time | Event | ms | 1,422 | 2,771 | 93 | | | | 1,158 | 4,001 | 61 | Kolmogorov-Smirnov | | 0,1964 | 0,1167 | ns |
| | **E** | Decay Time | Event | ms | 6,674 | 419,9 | 131 | | | | 4,135 | 22,69 | 76 | Kolmogorov-Smirnov | | 0,3496 | <0,0001 | **** |
| | **G** | Nb of isol. events | Cell | Nb | 88,7 | 46,73 | 10 | | | | 72,2 | 41,27 | 10 | Mann-Whitney,Two-tailed | | | 0,4136 | ns |
| | **I** | Frequency | Cell | Hz | 0,4785 | 0,83 | 10 | | | | 0,3375 | 0,631 | 10 | Mann-Whitney,Two-tailed | | 39,5 | 0,4475 | ns |
| | **J (left)** | Amplitude | Event | pA | 20,12 | 1286 | 887 | | | | 30,06 | 1409 | 722 | Kolmogorov-Smirnov | | 0,2776 | <0,0001 | **** |
| | **J (bottom left)** | Amplitude <50pA | Event | pA | 17,7 | 39,68 | 717 | | | | 25,63 | 39,69 | 574 | Kolmogorov-Smirnov | | 0,3381 | <0,0001 | **** |
| | **J (bottom right)** | Amplitude >50pA | Event | pA | 90,73 | 1246 | 170 | | | | 73,24 | 1369 | 148 | Kolmogorov-Smirnov | | 0,1955 | 0,0047 | ** |
| | **J (right)** | Decay Time | Event | ms | 5,514 | 46,22 | 887 | | | | 6,199 | 46,73 | 722 | Kolmogorov-Smirnov | | 0,08962 | 0,0033 | ** |
| | | | | | Mean | SD | N | | | | Mean | SD | N | Test | | Value test | P-value | |
| | **K** | Delta Decay | Paired events | | 8,2 | 7,9 | 10 | | | | | | | Wilcoxon signed rank test | | | 1.0 | ns |
| | | | Paired events | | | | | | | | -3,7 | 1,9 | 7 | Wilcoxon signed rank test | | | 0.047 | * |
| Supplemental data | | | | | | | | | | | | | | | | | | |
| Figure | Panel | Data | | | Control (iNGN) | | | | | | | | | Statisitics | | | | |
| **Suppl. Figure 1** | | | | | Mean | SD | N | | | | | | | Test | | Value test | Padj-value | |
| | **F** | PSD-95 density index | Dendrite | 7dpi | 0,03618 | 0,02129 | 36 | | | | | | | One-way ANOVA, Tukey's | 7 dpi vs. 14 dpi | F=52.18 | 0,0682 | ns |
| | | | | 14dpi | 0,0655 | 0,01655 | 11 | | | | | | | | 7 dpi vs. 34 dpi | | <0,0001 | **** |
| | | | | 34dpi | 0,08878 | 0,02925 | 13 | | | | | | | | 7 dpi vs. 56 dpi | | <0,0001 | **** |
| | | | | 56dpi | 0,1155 | 0,0334 | 18 | | | | | | | | 7 dpi vs. 72 dpi | | <0,0001 | **** |
| | | | | 72dpi | 0,1534 | 0,04799 | 23 | | | | | | | | 14 dpi vs. 34 dpi | | 0,3949 | ns |
| | | | | | | | | | | | | | | | 14 dpi vs. 56 dpi | | 0,0009 | *** |
| | | | | | | | | | | | | | | | 14 dpi vs. 72 dpi | | <0,0001 | **** |
| | | | | | | | | | | | | | | | 34 dpi vs. 56 dpi | | 0,1574 | ns |
| | | | | | | | | | | | | | | | 34 dpi vs. 72 dpi | | <0,0001 | **** |
| | | | | | | | | | | | | | | | 56 dpi vs. 72 dpi | | 0,0026 | ** |
| | **G** | vGlut1 puncta/neurite | Neurite | 7dpi | 0,36 | 0,2515 | 36 | | | | | | | One-way ANOVA, Tukey's | 7 dpi vs. 14 dpi | F=19.99 | 0,9793 | ns |
| | | | | 14dpi | 0,4145 | 0,1582 | 11 | | | | | | | | 7 dpi vs. 34 dpi | | 0,0361 | * |
| | | | | 34dpi | 0,6215 | 0,2299 | 13 | | | | | | | | 7 dpi vs. 56 dpi | | <0,0001 | **** |
| | | | | 56dpi | 1,018 | 0,3259 | 19 | | | | | | | | 7 dpi vs. 72 dpi | | <0,0001 | **** |
| | | | | 72dpi | 0,7346 | 0,3373 | 23 | | | | | | | | 14 dpi vs. 34 dpi | | 0,3707 | ns |
| | | | | | | | | | | | | | | | 14 dpi vs. 56 dpi | | <0,0001 | **** |
| | | | | | | | | | | | | | | | 14 dpi vs. 72 dpi | | 0,0187 | * |
| | | | | | | | | | | | | | | | 34 dpi vs. 56 dpi | | 0,0013 | ** |
| | | | | | | | | | | | | | | | 34 dpi vs. 72 dpi | | 0,7678 | ns |
| | | | | | | | | | | | | | | | 56 dpi vs. 72 dpi | | 0,012 | * |

Supplemental data

| Figure | Panel | Data | | | Control (iNGN) | | | | | | | | | Statisitics | | | | |
|---|---|---|---|---|---|---|---|---|---|---|---|---|---|---|---|---|---|---|
| **Suppl. Figure 1** | | | | | Mean | SD | N | | | | | | | Test | | Value test | Padj-value | |
| | **H** | vGlut positive synapses | Dendrite (%) | 7dpi | 27,64 | 11,55 | 36 | | | | | | | One-way ANOVA, Tukey's | 7 dpi vs. 14 dpi | F=16.47 | 0,0029 | ** |
| | | | | 14dpi | 43,93 | 10,8 | 11 | | | | | | | | 7 dpi vs. 34 dpi | | 0,9858 | ns |
| | | | | 34dpi | 29,75 | 11,93 | 13 | | | | | | | | 7 dpi vs. 56 dpi | | <0,0001 | **** |
| | | | | 56dpi | 54,67 | 17,97 | 18 | | | | | | | | 7 dpi vs. 72 dpi | | 0,0003 | *** |
| | | | | 72dpi | 42,46 | 10,45 | 23 | | | | | | | | 14 dpi vs. 34 dpi | | 0,0568 | ns |
| | | | | | | | | | | | | | | | 14 dpi vs. 56 dpi | | 0,1842 | ns |
| | | | | | | | | | | | | | | | 14 dpi vs. 72 dpi | | 0,9978 | ns |
| | | | | | | | | | | | | | | | 34 dpi vs. 56 dpi | | <0,0001 | **** |
| | | | | | | | | | | | | | | | 34 dpi vs. 72 dpi | | 0,0375 | * |
| | | | | | | | | | | | | | | | 56 dpi vs. 72 dpi | | 0,0236 | * |
| Figure | Panel | Data | | | Control-CRISPRa | | | | | | ERV-W-CRISPRa | | | Statisitics | | | | |
| **Suppl. Figure 3** | | | | | Mean | SD | N | | | | Mean | SD | N | Test | | Value test | P-value | |
| | **C** | Intencity | Cell | a.u. | 19,21 | 0,349 | 15 | | | | 19,42 | 0,3809 | 12 | Unpaired t-test, two-tailed | Cont.-ERVW | | 0,8799 | ns |
| Figure | Panel | Data | | | Control-CRISPRa | | | Syncytin-1-CRISPRa | | | ERV-W-CRISPRa | | | Statisitics | | | | |
| **Suppl. Figure 4** | | | | | Mean | SD | N | Mean | SD | N | Mean | SD | N | Test | | Value test | Padj-value | |
| | **B (top)** | Membrane capacitance | Cell | pF | 56,19 | 19,41 | 13 | 55 | 21,28 | 8 | 51,67 | 18,86 | 12 | One-way ANOVA, Bonferroni | Cont.-ERVW | F=3,02 | 0,5501 | ns |
| | | | | | | | | | | | | | | | Cont.-Syn. | | 0,6446 | ns |
| | | | | | | | | | | | | | | | ERVW-Syn. | | 0,0622 | ns |
| | **B (bottom)** | Input resistance | Cell | MΩ | 135,7 | 67,99 | 13 | 103,9 | 48,05 | 8 | 150 | 85,61 | 12 | One-way ANOVA, Bonferroni | Cont.-ERVW | F=3,02 | 0,5501 | ns |
| | | | | | | | | | | | | | | | Cont.-Syn. | | 0,6446 | ns |
| | | | | | | | | | | | | | | | ERVW-Syn. | | 0,0622 | ns |
| | **C** | Area | Cell | cm² | 790,223 | 197,1933333 | 1319 | 767,43 | 343,4625 | 642 | 761,777 | 314,4363333 | 1098 | One-way ANOVA, Bonferroni | Cont.-ERVW | | >0,9999 | ns |
| | **D** | Resting membrane potential | Cell | mV | -54,92 | 6,022 | 12 | -54,13 | 3,944 | 8 | -43,87 | 11,95 | 12 | One-way ANOVA, Bonferroni | Cont.-ERVW | F=6.052 | 0,01 | * |
| | | | | | | | | | | | | | | | Cont.-Syn. | | >0,9999 | ns |
| | | | | | | | | | | | | | | | ERVW-Syn. | | 0,0382 | * |
| | **F (top left)** | Fmax | Cell | Hz | 17,23 | 7,373 | 13 | 19,5 | 9,181 | 8 | 8,333 | 4,658 | 11 | One-way ANOVA, Bonferroni | Cont.-ERVW | F=9,259 | 0,0085 | ** |
| | | | | | | | | | | | | | | | Cont.-Syn. | | 0,7356 | ns |
| | | | | | | | | | | | | | | | ERVW-Syn. | | 0,0011 | ** |
| | **F (top right)** | Peak Amplitude | Cell | mV | 41,89 | 8,51 | 13 | 48,09 | 10,11 | 8 | 26,35 | 15,29 | 11 | One-way ANOVA, Bonferroni | Cont.-ERVW | F=9,259 | 0,0085 | ** |
| | | | | | | | | | | | | | | | Cont.-Syn. | | 0,7356 | ns |
| | | | | | | | | | | | | | | | ERVW-Syn. | | 0,0011 | ** |
| | **F (bottom left)** | Half width | Cell | msec | 2,567 | 0,6753 | 13 | 1,939 | 0,4719 | 8 | 4,588 | 1,878 | 11 | One-way ANOVA, Bonferroni | Cont.-ERVW | F=13,31 | 0,001 | *** |
| | | | | | | | | | | | | | | | Cont.-Syn. | | 0,7698 | ns |
| | | | | | | | | | | | | | | | ERVW-Syn. | | 0,0002 | *** |
| | **F (bottom right)** | Threshold | Cell | mV | -31,86 | 3,764 | 13 | -31,68 | 4,453 | 7 | -26,29 | 5,886 | 12 | One-way ANOVA, Bonferroni | Cont.-ERVW | F=4,927 | 0,0215 | * |
| | | | | | | | | | | | | | | | Cont.-Syn. | | >0,9999 | ns |
| | | | | | | | | | | | | | | | ERVW-Syn. | | 0,0756 | ns |
| | **H** | Sodium current peak | Cell | pA | -6763 | 3549 | 13 | -7939 | 3571 | 8 | -3616 | 2021 | 12 | One-way ANOVA, Bonferroni | Cont.-ERVW | | 0,0485 | * |
| | | | | | | | | | | | | | | | Cont.-Syn. | | >0,9999 | ns |
| | | | | | | | | | | | | | | | ERVW-Syn. | | 0,0135 | * |
| Figure | Panel | Data | | | Control-CRISPRa | | | Syncytin-1-CRISPRa | | | ERV-W-CRISPRa | | | Statisitics | | | | |
| **Suppl. Figure 6** | | | | | Mean | SD | N | | | | Mean | SD | N | Test | | Value test | P-value | |
| | **C** | Ratio AUC | Cell | | 0,5 | 0,3996 | 7 | | | | 0,2949 | 0,2717 | 6 | Unpaired t-test, one-tailed | Cont.-ERVW | | 0,1557 | ns |
